# The protein organization of a red blood cell

**DOI:** 10.1101/2021.12.10.472004

**Authors:** Wisath Sae-Lee, Caitlyn L. McCafferty, Eric J. Verbeke, Pierre C. Havugimana, Ophelia Papoulas, Claire D. McWhite, John R. Houser, Kim Vanuytsel, George J. Murphy, Kevin Drew, Andrew Emili, David W. Taylor, Edward M. Marcotte

## Abstract

Red blood cells (RBCs, erythrocytes) are the simplest primary human cells, lacking nuclei and major organelles, and instead employing about a thousand proteins to dynamically control cellular function and morphology in response to physiological cues. In this study, we defined a canonical RBC proteome and interactome using quantitative mass spectrometry and machine learning. Our data reveal an RBC interactome dominated by protein homeostasis, redox biology, cytoskeletal dynamics, and carbon metabolism. We validated protein complexes through electron microscopy and chemical crosslinking, and with these data, built 3D structural models of the ankyrin/Band 3/Band 4.2 complex that bridges the spectrin cytoskeleton to the RBC membrane. The model suggests spring-link compression of ankyrin may contribute to the characteristic RBC cell shape and flexibility. Taken together, our study provides an in-depth view of the global protein organization of human RBCs and serves as a comprehensive resource for future research.

## INTRODUCTION

Red blood cells (RBCs) fill a critical role by transporting oxygen and metabolic waste between the lungs and other cells and tissues. This role has been optimized by the unique features of RBCs, which lack nuclei, mitochondria, golgi, the endoplasmic reticulum, and most other major organelles so as to maximize oxygen carrying capacity. Nonetheless, RBCs respond actively to changing tissue environments and dynamically alter their cell shapes on short time scales in order to thread narrow capillary networks and splenic tissues, pointing to the critical importance of protein interactions, allostery, and post-translational modifications as the likely primary mechanisms to regulate RBC activities. As a result, mutations that disrupt these aforementioned processes lead to various blood disorders such as spherocytosis and hemolytic anemia.

RBCs have been studied extensively for decades and have provided us with essential basic information on cell physiology and molecular biology. However, a consensus on their complete proteome has not been reached, nor is it known how these proteins organize into the large multiprotein assemblies that support all RBC function in the absence of transcriptional and translational activity. While hemoglobin comprises the vast majority of RBC protein content (∼98%)(Goodman et al., 2007; Kabanova et al., 2009; Pasini et al., 2006), more than a thousand other distinct proteins are predicted to constitute the remaining 2% of protein, many of which are still largely uncharacterized. Since proteins rarely act alone, building a more complete picture of multiprotein assemblies is key to better understanding RBC functions and diseases, including cell shape control and the “shape-opathies” that result from its disruption.

Although techniques such as affinity purification mass spectrometry (AP-MS) and proximity labeling are available to study protein-protein interactions (PPIs) in other cell types, these approaches are not feasible in RBCs because of the lack of nuclei and transcription, and antibody-based approaches such as immunoprecipitation-mass spectrometry (IP-MS) are prohibitively expensive for large-scale screening. Therefore, we turned to another powerful technique to study protein complexes, co-fractionation mass spectrometry (CF-MS). CF-MS is a high-throughput technique that combines biochemical fractionations, protein mass spectrometry, and machine learning to characterize PPIs, in which the co-elution (co-fractionation) of stably-associated proteins through the course of distinct and orthogonal biochemical separations serves as evidence for the proteins’ physical association (Skinnider and Foster, 2021; Wan et al., 2015). As CF-MS requires neither antibodies nor recombinant epitope tagging of individual proteins, it is uniquely well-suited to study RBCs. The power of this technique comes from the analysis of native proteins’ elution profiles from multiple orthogonal biochemical separations, using machine learning and extensive internal control proteins (known complexes) to distinguish between true PPIs and randomly co-eluting proteins, quantifying the support for the physiologically relevant PPIs in order to maintain strong control over false discovery rates of protein interactions.

In this work, we performed over 30 biochemical fractionations on purified RBCs and applied quantitative mass spectrometry on these fractions (>1,900 mass spectrometry experiments in all) in order to identify a canonical set of approximately 1,200 RBC proteins and to derive an interactome map of RBC protein complexes. Using a rigorous statistical framework, we recovered high confidence PPIs from known complexes as well as novel complexes. Most of the complexes in RBCs are involved in energy metabolism, structural integrity, redox biology, and proteostasis. Furthermore, we performed chemical crosslinking on these native complexes and used integrative structural modeling to shed light on the molecular organization of integral membrane proteins, channels, cytoskeletal proteins, and metabolic enzymes at the RBC membrane. Our findings provide a comprehensive blueprint of the cell surface and subcellular architecture of a key blood cell type, and suggest biophysical mechanisms underlying its adaptability and pathological reorganization in diverse blood disorders.

## RESULTS AND DISCUSSION

### A large data set of protein abundances and co-purification of RBC proteins

In order to characterize the RBC proteins and their assemblies, we generated a large proteomics dataset of fractionated soluble and membrane-associated proteins from enriched human RBCs using various methods of biochemical fractionation (**Figure 2A**). Non-denatured protein extracts from hemolysate (soluble proteins) and non-ionic detergent dissolved ghosts (membrane proteins) were separated by discrete biophysical properties such as size, charge, and hydrophobicity (see Zenodo data repository for details of fractionations, donors’ details, detergents used). Each chromatographic fraction was analyzed by high resolution, high sensitivity liquid chromatography/mass spectrometry (LC/MS). In all, we collected 6,255,027 interpretable peptide mass spectra from 1,944 individual chromatographic fractions. Each fraction captures distinct subsets of native RBC proteins and protein assemblies, collectively providing a compendium of informative co-elution profiles.

### Determination of a high-confidence RBC proteome

From this large dataset, we first aimed to comprehensively define a canonical set of RBC proteins. Although multiple previous surveys have identified versions of the RBC proteome (Bryk and Wiśniewski, 2017; Goodman et al., 2007; Lange et al., 2014), these studies only agree on 859 constituent proteins (**Figure 1A**). Discrepancies might arise from technical variation in the analyses and samples as well as from the presence of contaminating proteins contributed by accompanying reticulocytes, platelets, white blood cells (WBCs), and serum proteins. We attempted to reconcile these published datasets and our own data using a bioinformatic approach: We trained a machine learning classifier on available MS and RNA-seq data from different blood cell types (**Figure 1A**) in order to recognize the best supported consensus set of RBC proteins. The rationale for this approach is that some proteins detected in previous studies or our fractionation experiments inevitably derive from non-RBC blood cells despite best efforts to enrich RBCs. This issue is exacerbated by the fact that all non-hemoglobin proteins only represent 2% of the RBC proteins, and thus even 2% of contaminant cells could skew protein identifications for RBC proteome. The consideration of data spanning multiple cell types should allow RBC proteins to be better distinguished from proteins from other blood cells and serum.

**Figure 1.**
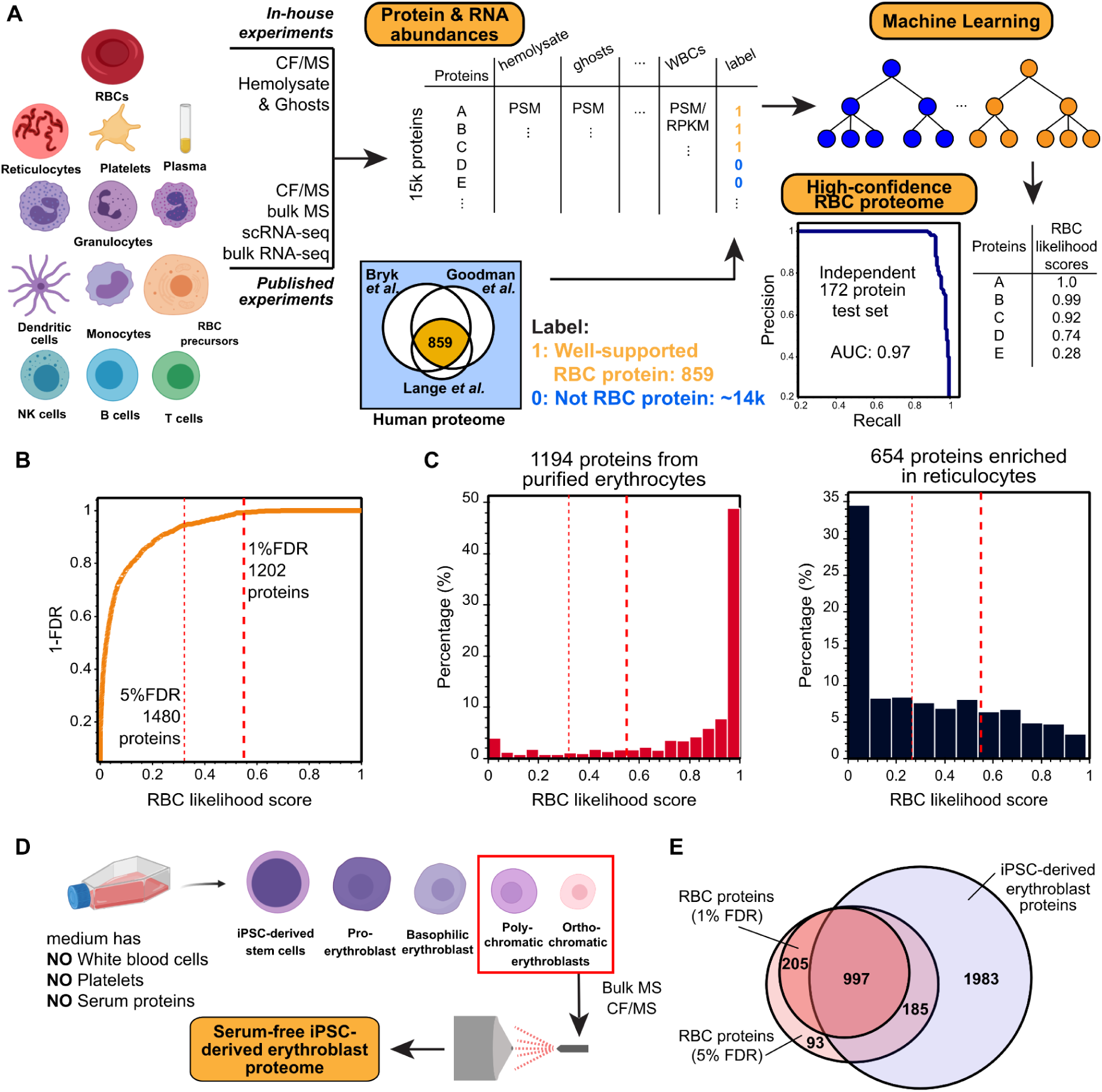
Defining the canonical red blood cell (RBC) proteome with high accuracy from a synthesis of protein mass spectrometry and mRNA expression data. **(A)** Measured protein and RNA abundances from diverse blood cell types and plasma were used as features for a machine learning classifier to assign confidence scores for proteins whether they belong to the RBCs or rather contaminants contributed by other cells or plasma. The classifier showed high precision and recall (area under the recall-precision curve (AUC) = 0.97) as assessed on a 172 protein set withheld from the training. Cell images created with Biorender.com. **(B)** Applying the classifier and thresholding at a 1% false discovery rate (FDR), we observe the canonical RBC proteome to comprise 1,202 proteins. **(C)** The resulting high confidence RBC proteins are highly concordant with proteins previously identified from purified erythrocytes (Gautier et al., 2018). In contrast, the 1% FDR proteome notably excludes proteins known to be strongly enriched in reticulocytes (Gautier et al., 2018). **(D)** To further assess the potential for proteins to be contributed from other blood cells or serum, we differentiated iPSC cells into polychromatic and orthochromatic erythroblasts in serum-free medium lacking white blood cells, platelets, and serum proteins, then analyzed the erythroblast proteome using mass spectrometry. **(E)** A large majority of high confidence RBC proteins at both the 1% and 5% FDR level (1,182 proteins in total) were also detectable in erythroblasts, consistent with our expectation that mature RBC proteins should generally be detectable in a relevant precursor cell population.

There are two major issues to consider when trying to identify the comprehensive list of RBC proteins: coverage and contamination. In terms of coverage, we detected >90% of the highly-confident set of 859 previously described proteins (**Figure 1**) among the 2,000 most abundant proteins from our experiments (**Figure S1A**). This analysis shows that our co-fractionation experiments were well-powered to detect most RBC proteins. Some exceptions are notable, particularly alpha and beta actins (ACTA, ACTB), which are systematically depleted due to our use of mild and non-ionic detergents to extract proteins while preserving stable interactions, and the Duffy antigen ACKR1.

In order to distinguish bona-fide RBC proteins from proteins from other cell types, we employed a supervised machine learning approach to quantify the likelihood of proteins deriving from RBCs. We analyzed protein and RNA abundances (from MS and RNA-seq data) from RBCs, RBC precursor cells, other blood cell types (including white blood cells, platelets, and reticulocytes), and serum (**Figure 1A**), with the derived likelihood score based solely on these measurements. As confirmed positive training examples, we selected a set of 687 known RBC proteins out of the 859 proteins agreed upon by all three reference studies, and we withheld the remaining 172 positive examples as an independent test set. As confirmed negative training examples, we considered all human proteins observed in our proteomics experiments that were not observed as RBC proteins by the 3 prior studies. We then trained a random forest classifier (with 5-fold cross-validation) to assign a confidence score between 0 and 1 to each protein (see Zenodo data repository for scored proteins), with 1 indicating a high likelihood of the protein existing in mature RBCs, and 0 indicating a likely contaminant.

We assessed the quality of these confidence scores by comparison to the withheld test set of 172 known RBC protein markers and a set of withheld negative test examples. The classifier performed extremely well as indicated by a high area under the precision-recall curve (AUPR) of 0.97 (**Figure 1A**). Ranking proteins based on their likelihood scores enabled us to measure classifier false discovery rates (FDR) (**Figure 1B**). At a likelihood score >0.55, we observed 1,202 proteins at 1% FDR (**Figure S1B**). We consider this stringent threshold to define a comprehensive and high confidence set of RBC proteins. Ranking the proteins detected in each of the three prior RBC proteome studies by our RBC likelihood score shows that our classifier efficiently distinguished RBC proteins from non-RBC proteins (**Figure S1C-E**). Among the reference studies, the preparation by Lange *et al*. comprised a notably high proportion of mature RBCs.

Besides well-established RBC proteins such as hemoglobin, carbonic anhydrase, RBC-specific membrane proteins (e.g., Band 3, Ankyrin 1, Band 4.2), and blood-type antigens (e.g., Rh antigens, Kell), the high-confidence RBC proteome faithfully reflects the unique biology of mature RBCs in that they lack major organelles, or the apparatus for DNA replication, transcription, and protein synthesis. To further support the proteome, because most proteins in mature RBCs would be already present in RBC precursor cells, we asked if they were additionally present in induced pluripotent stem cell (iPSC)-derived erythroblasts. While nucleated, these RBC precursors were grown in serum-free medium (Vanuytsel et al., 2018) and were devoid of all contaminant proteins from other blood cell types and from serum (**Figure 1D**). In all, 83% of the high-confidence RBC proteins in our compendium (1% FDR) were also present in the erythroblasts. Similarly, we compared the high confidence set of RBC proteins to those obtained through multi-step RBC purification, including centrifugation, gradient/cellulose purification, and fluorescence activated cell sorting (FACS), by Gautier and colleagues (Gautier et al., 2018). The majority of RBC-specific proteins obtained in this way exhibited high RBC likelihood scores, while the majority of proteins obtained from similarly enriched reticulocytes had markedly lower RBC likelihood scores (**Figure 1C**). Indeed, the high-confidence RBC proteome excluded CD71 (Transferrin receptor protein 1, TFRC), a well-known reticulocyte marker protein (Gautier et al., 2016, 2018; Goodman et al., 2007). Taken together, these results suggest that our bioinformatic approach effectively captured the proteome of mature RBCs and excluded most major contaminating non-RBC proteins.

Interestingly, some proteins with high RBC likelihood scores consisted of annotated subunits of ribosomes, nuclear pore assemblies, and parts of the ER and Golgi machineries, despite RBCs lacking these larger systems. Many of these proteins were also detected as RBC-specific populations by Gautier et al., 2018, and were observed in the iPSC-derived erythroblasts. Thus, they are unlikely to be contaminants. Although we can not rule out the possibility of moonlighting functions in RBCs, another explanation may simply be that such proteins are not eliminated or degraded effectively during the process of enucleation in reticulocytes, in which the nucleus was expelled, major organelles were eliminated, and other superfluous proteins were degraded. Thus, the proteome could at least in part reflect an imperfect cell maturation process.

### Systematic identification and scoring of stable RBC protein interactions

Given this high confidence set of RBC proteins, we next sought to determine their association into higher order multi-protein assemblies. In CF-MS, subunits of many well-known complexes coelute with distinct patterns across different types of biochemical separations due to differences in complex sizes, charges, and chemical properties (**Figure 2C**). To analyze such differences and identify co-eluting proteins in a systematic and high-throughput manner, we again used a supervised machine learning approach in order to assign a confidence score to each potential protein-protein interaction based on the mass spectrometry evidence (**Figure 2B**). Importantly, PPIs were derived solely from the separation behavior of the observed proteins over multiple, orthogonal biochemical fractionation experiments directly from RBCs, with no contribution from external data (see Methods and Zenodo data repository). We focused on abundant RBC proteins with the most robust mass spectrometry evidence (more than 60 independent peptide-spectral matches (PSMs) across 1,944 biochemical fractions). We trained an extra tree classifier (using 5-fold cross-validation) to distinguish between pairs of proteins known to stably interact and random pairs of proteins (details in Methods). The classifier assigned a probabilistic CF-MS score between 0 and 1 to each potential PPI (**Figure 2A**), with 1 indicating a high likelihood of physical association based on observing strongly coordinated protein elution profiles, and 0 indicating no evidence for interaction in RBCs.

**Figure 2.**
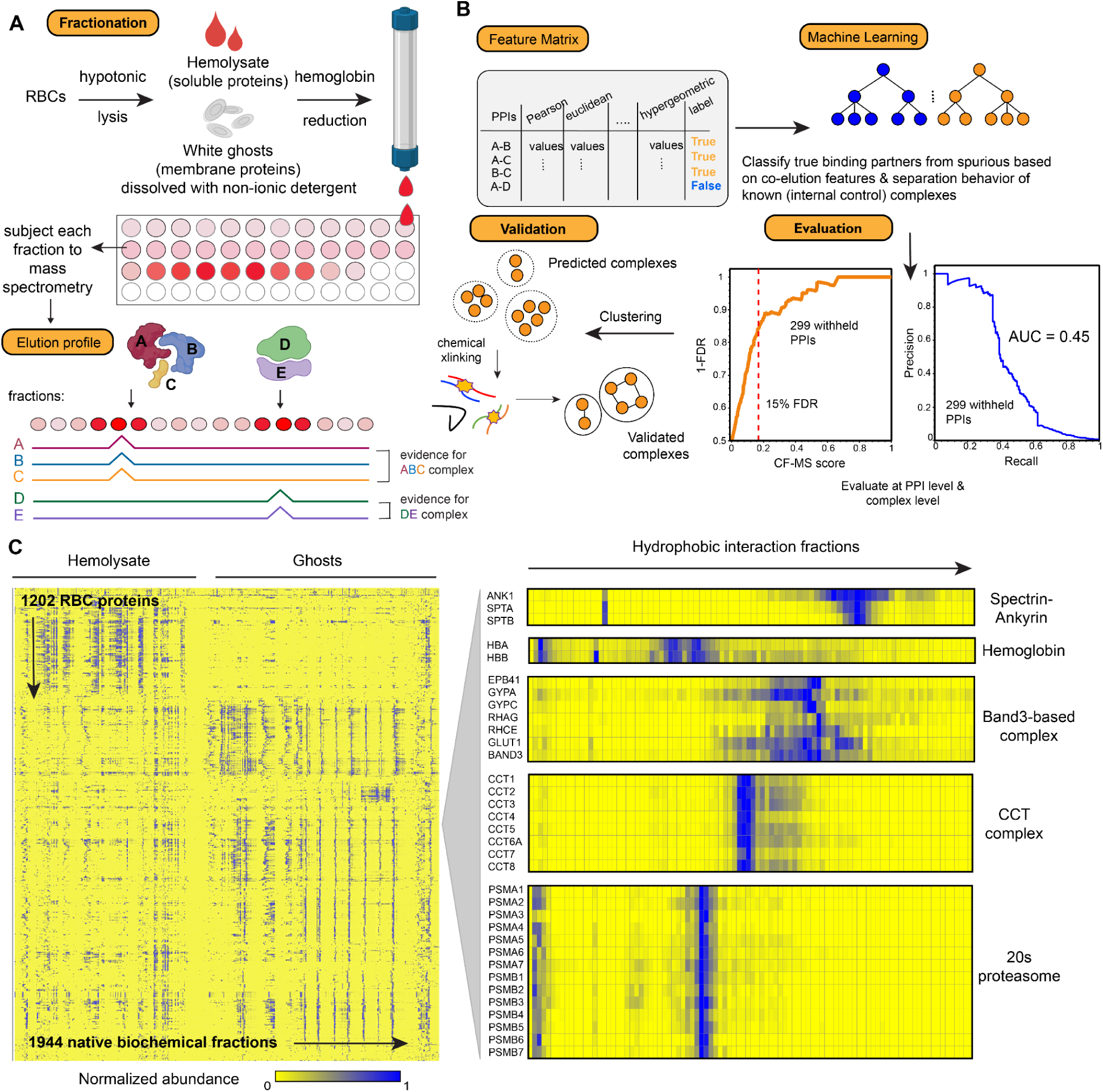
Overview of the integrative Co-Fractionation / Mass Spectrometry (CF-MS) workflow used to determine stable RBC protein complexes. **(A)** Hemolysate and white ghosts are chromatographically separated and the proteins in each fraction are identified by mass spectrometry. Elution profiles for each protein are graphically represented as ridgelines across multiple separation experiments. **(B)** Different measures of correlation between each pair of proteins are used to construct a feature matrix for machine learning, which computes a score (CF-MS score) indicating how likely the interaction between two proteins would be in RBCs. **(C)** Heat map of the full dataset of abundance measurements for each of the 1,202 RBC proteins across all fractionations of hemolysate and white ghosts. Blue indicates non-zero signal. **(D)** Enlarged portions of **(C)** showing examples of strong co-elution observed for subunits (gene names on left) of five well-known protein complexes in RBCs (complex names on right). Color intensity (blue is positive signal) depicts abundances for each protein from a representative hydrophobic interaction chromatography experiment (labeled on top) out of the 30 total separations.

We judged the quality of the measured interactions by comparing to a withheld test set of 235 known protein interactions (**Figure 2A**), which allowed us to measure classifier error rates. The CF-MS scores accurately recapitulated the test PPIs: for interactions with CF-MS scores over 0.17, we observed 85% precision and 35% recall (**Figure S3**). In all, we observed 3229 PPIs among the high-confidence RBC proteins that scored 0.17 or better, providing an estimate of the RBC interactome.

### Direct visualization of biochemically fractionated complexes by electron microscopy

Because CF-MS involves biochemical enrichment and separation of intact endogenous protein complexes, we could use electron microscopy (EM) to directly visualize the larger complexes in a relatively unbiased fashion (Kastritis et al., 2017; Verbeke et al., 2018). First, to survey the size, shape, and complexity of assemblies across a representative biochemical separation, we performed negative stain EM on pooled, adjacent fractions from a size exclusion chromatography separation (**Figure 3A**). By comparing the micrographs with proteins identified in the corresponding mass spectrometry experiments and prior knowledge of 3D structures, we identified four distinct protein complexes across the fractions (**Figure 3B**), three of which were large homo-oligomeric assemblies. The Tripeptidyl-peptidase 2 (TPP2) was the largest observed homo-oligomer and is known to form 5-6 MDa assemblies (Macpherson et al., 1987; Schönegge et al., 2012). The other homo-oligomers identified were the porphyrin-biosynthetic enzyme delta-aminolevulinic acid dehydratase (ALAD) (Mills-Davies et al., 2017), an ∼290 kDa homo octamer with D4 symmetry, and the peroxiredoxin PRDX2 (Schröder et al., 2000), an ∼218 kDa homo decamer with D5 symmetry.

**Figure 3.**
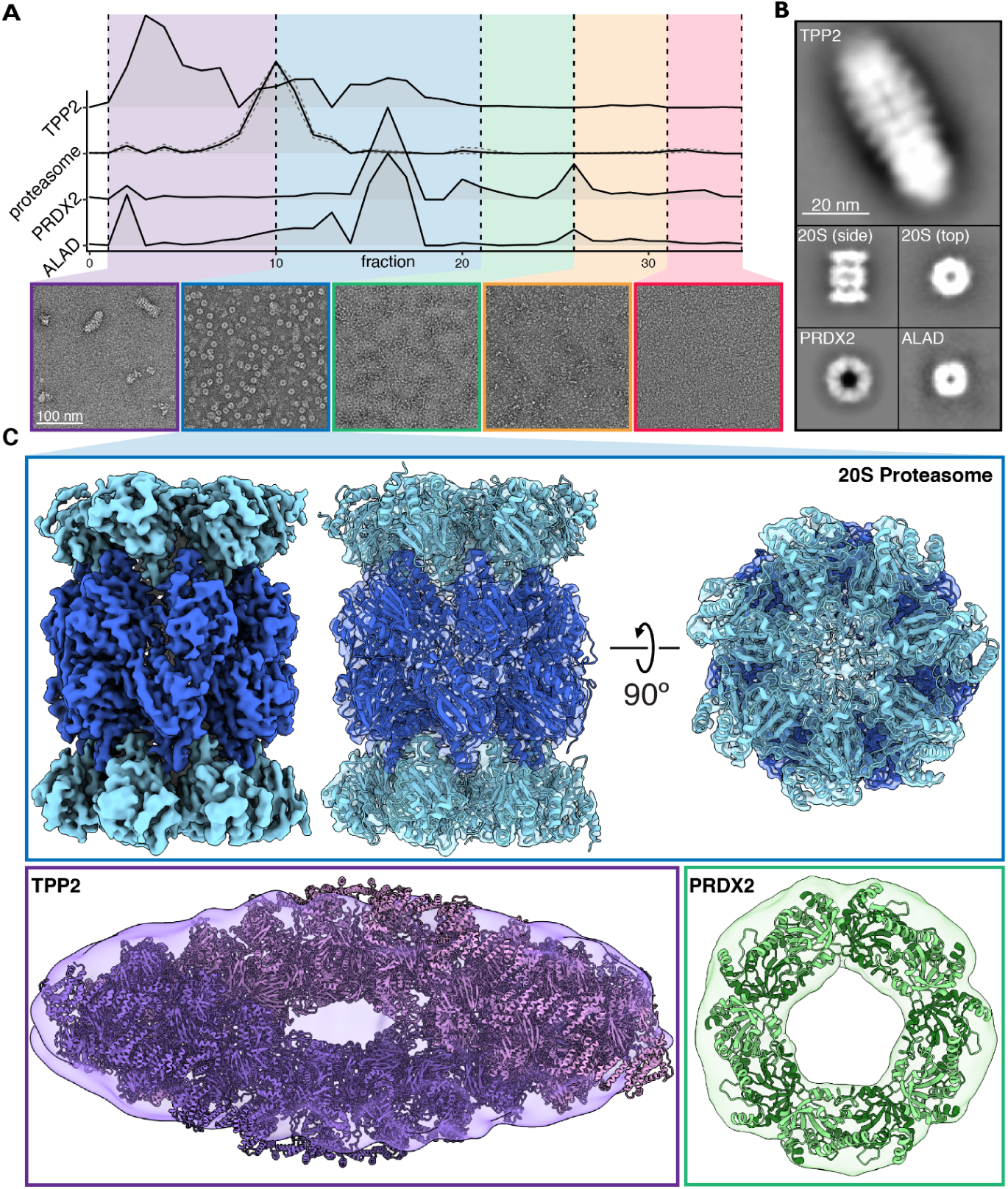
Validation of the CF-MS workflow using electron microscopy confirms intact multi-protein complexes. (**A**) Hemolysate from size exclusion chromatography was partitioned into five groups and visualized with negative stain EM. Elution profiles from corresponding mass spectrometry data were used to assist in identifying abundant protein assemblies. (**B**) Reference-free 2D class averages of four protein complexes spanning ∼220 – 5,000 kDa were identified in hemolysate. (**C**) Cryo-EM reconstruction of the 20S proteasome. Negative stain structures of TPP2 and PRDX2 along with docking of their corresponding atomic structures PDB 3LXU (Chuang et al., 2010) and PDB 1QMV (Schröder et al., 2000), respectively.

These data also clearly showed a high abundance of proteasomes, which provided an opportunity to revisit an outstanding question as to whether proteasomes in mature RBCs are active (*i.e*., in their regulated 26S forms) or mostly inactive (with separated 20S cores and 19S caps). We first pooled the proteasome containing fractions (fractions 10-21) for higher resolution analysis by cryo-EM. By single particle analysis, we obtained a reconstruction of the 20S proteasome directly from RBCs with a nominal resolution of 3.35 Å, confirming an intact, not visibly modified core proteasome (**Figure 3C, Figure S2**). To further investigate the proteasome’s activity in RBCs, we used negative stain EM to survey hemolysate after being passed through a 100 kDa filter and quantified different assembly states of the proteasome (**Figure S3**). We found that ∼94% of the proteasomes observed were of the free 20S form, while the remaining ∼6% were singly-capped 26S proteasomes (**Figure S3**). These results agreed with our CF-MS observations of generally separated 19S and 20S proteasomes, and suggested that active 26S proteasomes were present only in relatively low abundances.

### The interactome of mature red blood cells

The visualization of known complexes by electron microscopy confirmed that stable multiprotein assemblies were preserved well across our biochemical separations, so we next sought to systematically define protein complexes in mature RBCs by clustering the proteins based on the measured pairwise interactions. To provide a more comprehensive view of protein complexes, we elected to use multiple clustering cutoffs to reflect the hierarchies intrinsic to interacting proteins, the results of which are visualized in **Figure 4**.

**Figure 4.**
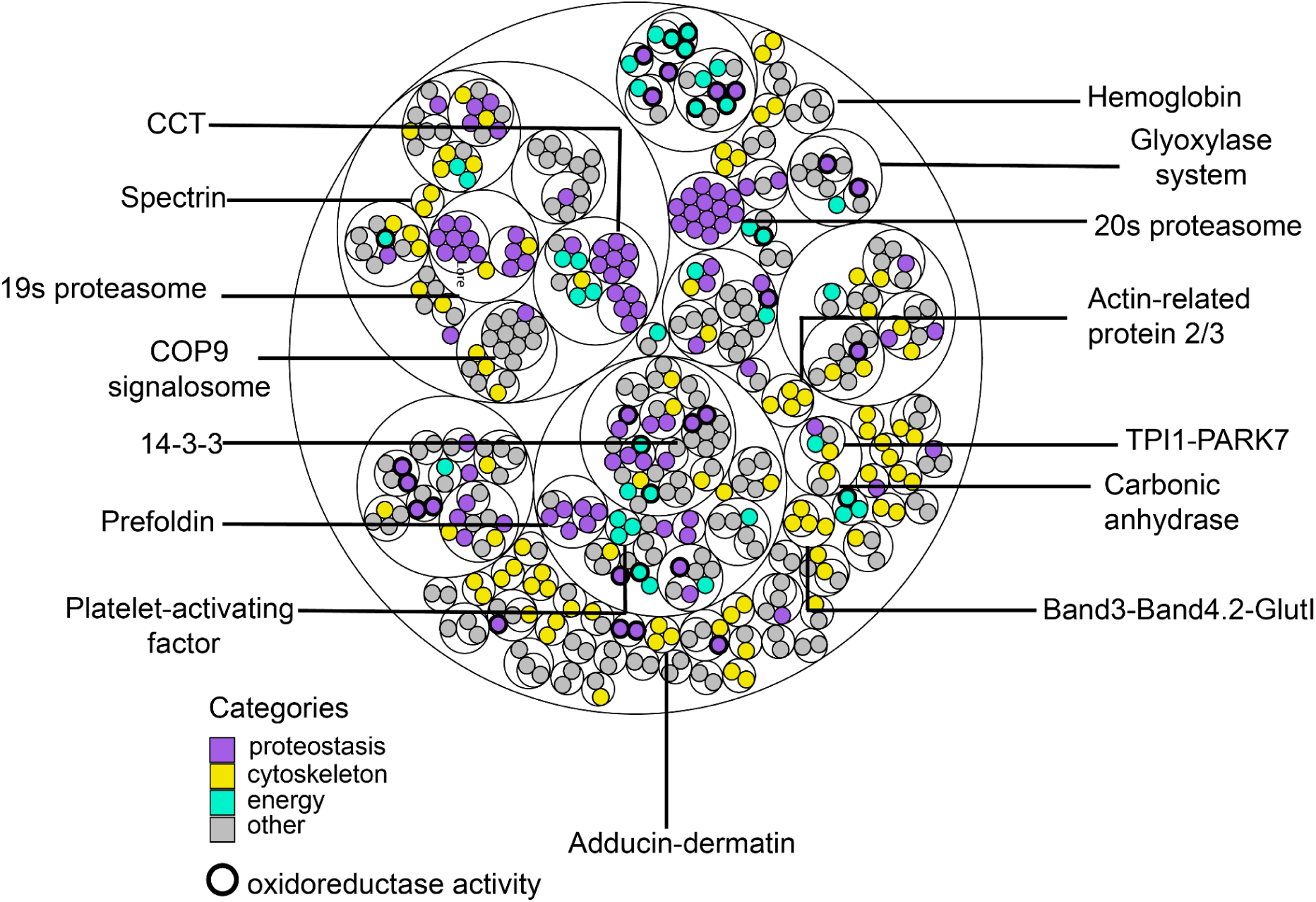
A map of the primary RBC multiprotein complexes. Thin circles show the clustering hierarchy of protein-protein interactions into complexes for each of 3 clustering thresholds (see the Zenodo data repository for complex memberships and annotations). Proteins (filled circles) are colored by broad biological categories (analyzed using the DAVID annotation tool); bold outlines denote proteins with oxidoreductase activity.

In all, the interactome of mature RBCs represents a remarkably compact and minimal assembly of protein machinery. First, we find just over 100 protein complexes in mature RBCs. While this is already apparent at the proteome level (∼1,200 total proteins in mature RBCs, or approximately ¼ the complexity of the proteome of the bacterium *E. coli* and 1/16 of the consensus human proteome), the number of complexes is still striking in comparison to the >400 complexes in *E. coli* (Hu et al., 2009) and >7,000 human protein complexes known to date (Drew et al., 2021). To gain some insights into the interactome, we analyzed the proteins according to their broad functional categories (Huang et al., 2009a, 2009b), and found most categorized proteins mapped into 4 major categories: proteostasis (purple), cytoskeleton/structural integrity (yellow), energy (mint green), and others (grey).

Proteins of proteostasis dominate the observed RBC interactome, notable among these being the 19S regulatory cap and 20S core of the proteasome. However, consistent with the evidence from EM micrographs (**Figure S3**), the clustering of PPIs suggests that proteasomes in mature RBC generally exist as distinct 19S and 20S forms. Notably, as there is no protein synthesis, and the lifetime of RBCs is only ∼120 days, protein degradation may be kept to minimal levels in mature RBCs. In contrast, we observe intact protein folding and aggregation dissociation chaperones, including the prefoldin and CCT chaperones.

Besides proteostasis, energy production and redox-related protein complexes were abundant. A survey of the interactome suggests a balance in regulating energy production and managing toxic by-products through detoxification and redox enzymes (bolded circle, **Figure 4**). RBCs consume energy to maintain proper ion concentrations and appropriate ratios of surface area to cell volume (Lux, 2016; Mohandas and Gallagher, 2008), which are essential to the ability of RBCs to change morphology without lysing. The major source of ATP production in RBCs involves the metabolism of glucose *via* the Embden-Myerhoff or glycolysis pathway (Brown, 1996). One of the toxic byproducts is methylglyoxal, a highly reactive dicarbonyl compound that reacts with proteins through the Maillard reaction to form glycated proteins, impairing protein function. In order to detoxify methylglyoxal, cells rely on the glyoxalase system/complex, which is recapitulated in the interactome. We find evidence for a novel interaction relevant to methylglyoxal detoxification occurring between triosephosphate isomerase (TPI1) and the Parkinsonism-associated protein (DJ-1/Park7). TPI1 is known to produce a significant amount of methylglyoxal as a byproduct (Richard., 1991), and Park7 is proposed to be a bonafide deglycase (Richarme & Laidrou., 2017). This TPI1-Park7 interaction suggests a physical linkage between energy production and byproduct detoxification in RBCs.

Finally, cytoskeletal complexes are essential to RBCs, as they maintain structural integrity and enable cell morphology changes as RBCs traverse blood vessels. The interactome recapitulates well-known cytoskeletal complexes involving, among other proteins, spectrin, adducin-dermatin, and band 3, and suggests subcomplexes such as Band 3, GlutI, and Band 4.2, as described in more detail below.

### Cross-linking and integrative 3D modeling of the major RBC cytoskeletal complexes

As cytoskeletal complexes are essential to many RBC functions, we particularly sought to investigate protein complexes isolated from RBC white ghosts, which are composed almost exclusively of membrane/cytoskeletal components. Thus, we separated detergent-solubilized white ghost protein extracts by size exclusion chromatography and performed chemical crosslinking mass spectrometry (XL-MS) in order to precisely determine amino acid-resolution PPI contacts between membrane and cytoskeletal proteins. We chose disuccinimidyl sulfoxide (DSSO) for crosslinking, as it is MS-cleaveable and thus suitable for highly accurate determination of crosslinked peptides (Kao et al., 2011). In all, we identified 769 high confidence crosslinks among 129 proteins, dominated by interactions among spectrins, ankyrin, and their interaction partners, including the Rh antigens, Band 3, and Band 4.1 (**Figure 5A**). Crosslinked proteins were more likely to have high CF-MS scores, suggesting that the experiment captured true molecular interactions (**Figure S4**). In addition to intermolecular crosslinks (**Figure 5A**), we observed large numbers of intramolecular crosslinks, as highlighted in **Figure 5B** for spectrin alpha and beta.

**Figure 5.**
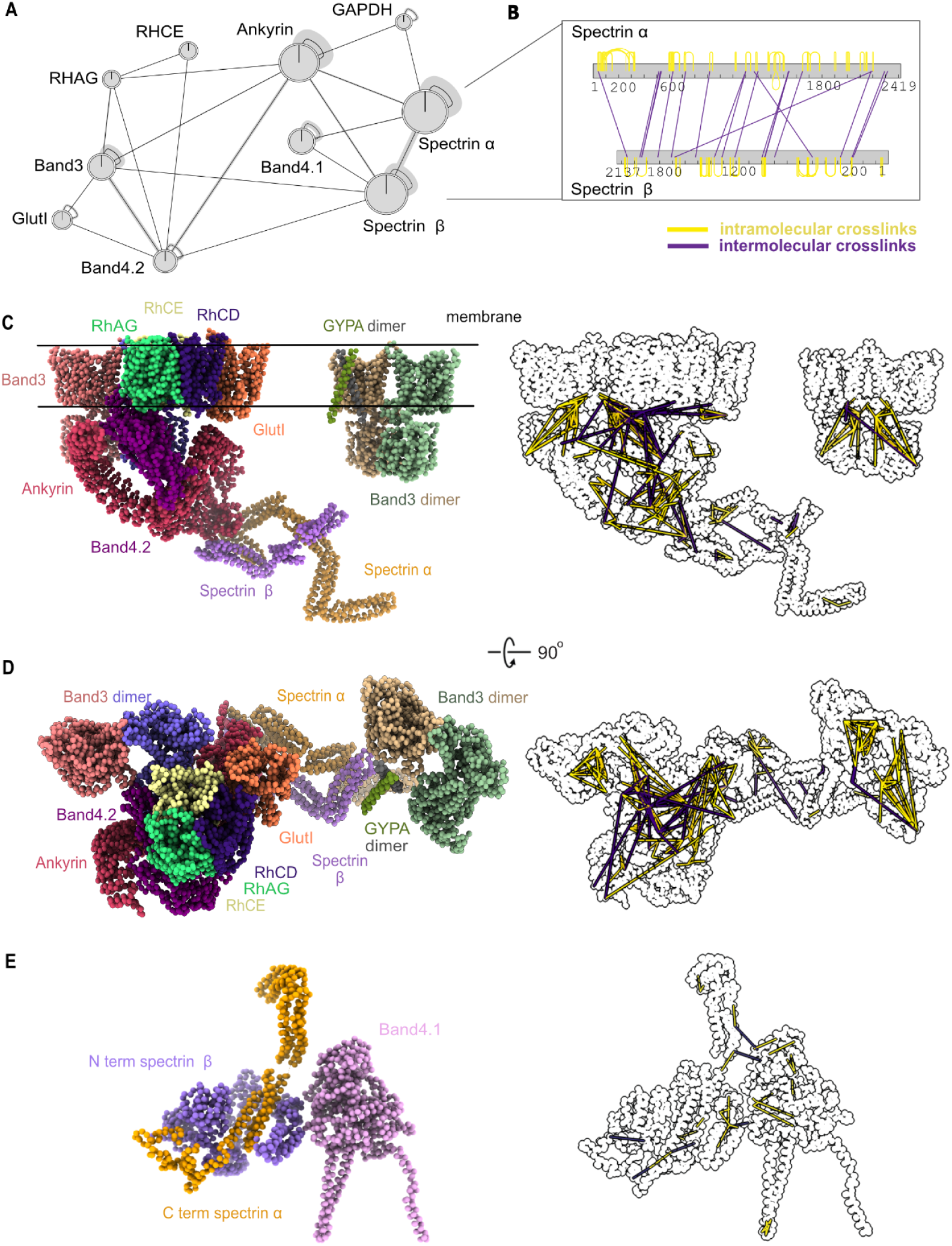
Chemical crosslinks confirm mapped interactions and constrain 3D modeling of membrane/cytoskeletal complexes. (A) Network plot represents crosslinked interactions among genes in RBC cytoskeletal complexes. Line between each node indicates detected crosslink(s). Shaded lines indicate dense crosslinking. (B) Bar plot shows extensive crosslinks between spectrin alpha and beta supporting a head-to-tail conformation. Numbers under each bar indicate the amino acid position on each protein. Yellow line indicates intramolecular crosslinks and purple line indicates intermolecular crosslinks. (C) Side view of integrative structure of band 3-Ank1 complex and band 3-GYPA complex. Our model suggests that GlutI competes with the Rh proteins for binding with band 3 and band 4.2. The outline figure on the right shows intramolecular and intermolecular crosslinks that are overlaid onto the structure. (D) Top view of integrative structure of the band 3-Ank1 complex and band 3-GYPA complex. (E) Integrative structure of band 4.1-spectrin complex. Six of these complexes are proposed to link spectrin heterodimers with actin (Lux, 2016).

Each crosslink generally provides evidence for the modified lysines residing within 30 Å of each other. Thus, such data enable initial molecular models to be constructed of the component proteins. For example, the pattern of crosslinks between spectrin alpha and beta strongly supports them associating in a head-to-tail conformation (**Figure 5B**), consistent with literature reports that the N-terminal domain of spectrin alpha interacts with the C-terminus of spectrin beta (Speicher et al., 1992; Ungewickell and Gratzer, 1978). To further evaluate the quality of the crosslinks, we used known X-ray crystal structures of the proteins band 3 (PDB ID: 1HYN, 4YZF), GAPDH (PDB ID: 1U8F), and PGK1 (PDB ID: 4O33) to calibrate the crosslinks’ accuracies. Of the 24 crosslinks between residue pairs within these X-ray crystal structures, all 24 occurred between amino acids with Cα atoms falling within 30 Å of each other, consistent with the length of the DSSO crosslinker (**Figure S4**).

Next, we integrated the crosslinks with available structural information in order to build models of several of the major RBC membrane/cytoskeletal complexes. We additionally incorporated knowledge of the positions of transmembrane regions (The UniProt Consortium, 2019), protein structures from comparative modeling, and, where available, X-ray crystal structures. Using integrative modeling (**Figure S5**) (Russel et al., 2012), we built many (600,000) models in parallel and tested for their agreement (see Methods); the combination of annotated transmembrane regions, chemical crosslinks, and known partial structures provided sufficient spatial restraints to converge on a solution. High-scoring models that satisfied the crosslinking and membrane restraints were clustered based on root-mean-square distance (RMSD), and the largest cluster (**Table S2**) was selected as the preferred form of each complex, with the centroid model as the representative model for the cluster.

This integrative modeling approach provided us with a detailed molecular view of both the band 3 (**Figure 5C-D**) and band 4.1 complexes (**Figure 5E**).Yellow and purple lines to the right of the models in **Figure 5C-E** highlight positions of intramolecular crosslinks and intermolecular crosslinks, respectively. Surprisingly, 30% of the crosslinks for ankyrin1 violated the 30 Å distance of the crosslinker, far in excess of the typically fewer than 20% of violations expected in such models (Leitner et al., 2016; Liu et al., 2018; Wang et al., 2017) and the 100% concordance we observed with the isolated band 3 and PGK1 structures, and suggesting that the reported X-ray crystal structure of ankyrin1 (Wang et al., 2014) does not match the predominant form found in RBCs. Notably, the reported structure is in an extended (“open”) form. Our crosslinks suggest that the violation could be due to ankyrin adopting a compressed (“closed”) conformation, as the violations are highly directionally correlated in two specific regions of ankyrin (**Figure S6**). When modeling a closed conformation of the ankyrin complex, 92% of the crosslink violations for ankyrin1 can be satisfied (**Figure S6**). We speculate that the open and closed conformations could be a result of spring-like behavior of ankyrin repeats; indeed, spring-like behavior has been previously observed for ankyrin by atomic force microscopy (Lee et al., 2006). Given ankyrin’s privileged position connecting the RBC plasma membrane (*via* interactions with the band 3/Rh membrane proteins) to the spectrin cytoskeleton, it is plausible that spring-like behavior of ankyrin may play an important role in maintaining RBCs’ morphologies as they deform while traversing microvasculature and splenic tissues.

Indeed, the band 3-ankyrin complex has been proposed to locate in the middle of the spectrin tetramer chain to anchor spectrins to the plasma membrane, while the band 4.1 complex connects the end of the tetrameric spectrin chain to one of six binding sites on actin in the actin junctional complex (Lux, 2016; Mohandas and Gallagher, 2008) (**Figure 6A**). Our model of the band 3-ankyrin complex shows the association of band 3 with band 4.2, Rh proteins, GlutI, ankyrin, spectrin and glycophorin A. The association of band 3 with band 4.2 and GlutI based on crosslinking correspond well with the cluster derived from CF-MS (**Figure 4**). In addition, our model of the band 3-ankyrin complex is supported by previous co-immunoprecipitation data from RBCs of an individual with almost complete absence of band 3 (Coimbra mutation, homozygous null for band 3), crystallography, *in vitro* binding assays, and mass spectrometry (Bruce et al., 2003; Ipsaro and Mondragón, 2010; Ipsaro et al., 2009; Jiang et al., 2006; Kumpornsin et al., 2011; Lemmon et al., 1992). A previously observed reduction of Rh proteins and band 4.2 in the absence of band 3 suggests that they also form a complex in RBCs (Bruce et al., 2003). Interestingly, Bruce and colleagues found that GlutI was detected more in the absence of band 3. Our cross-linking evidence and 3D model indicates that GlutI competes with Rh proteins at the same binding sites/regions on band 3 and band 4.2, while band 3 and glycophorin A form a separate complex. These data thus support at least 3 subpopulations of band 3 complexes: those with Rh proteins, GlutI, or glycophorin A.

**Figure 6.**
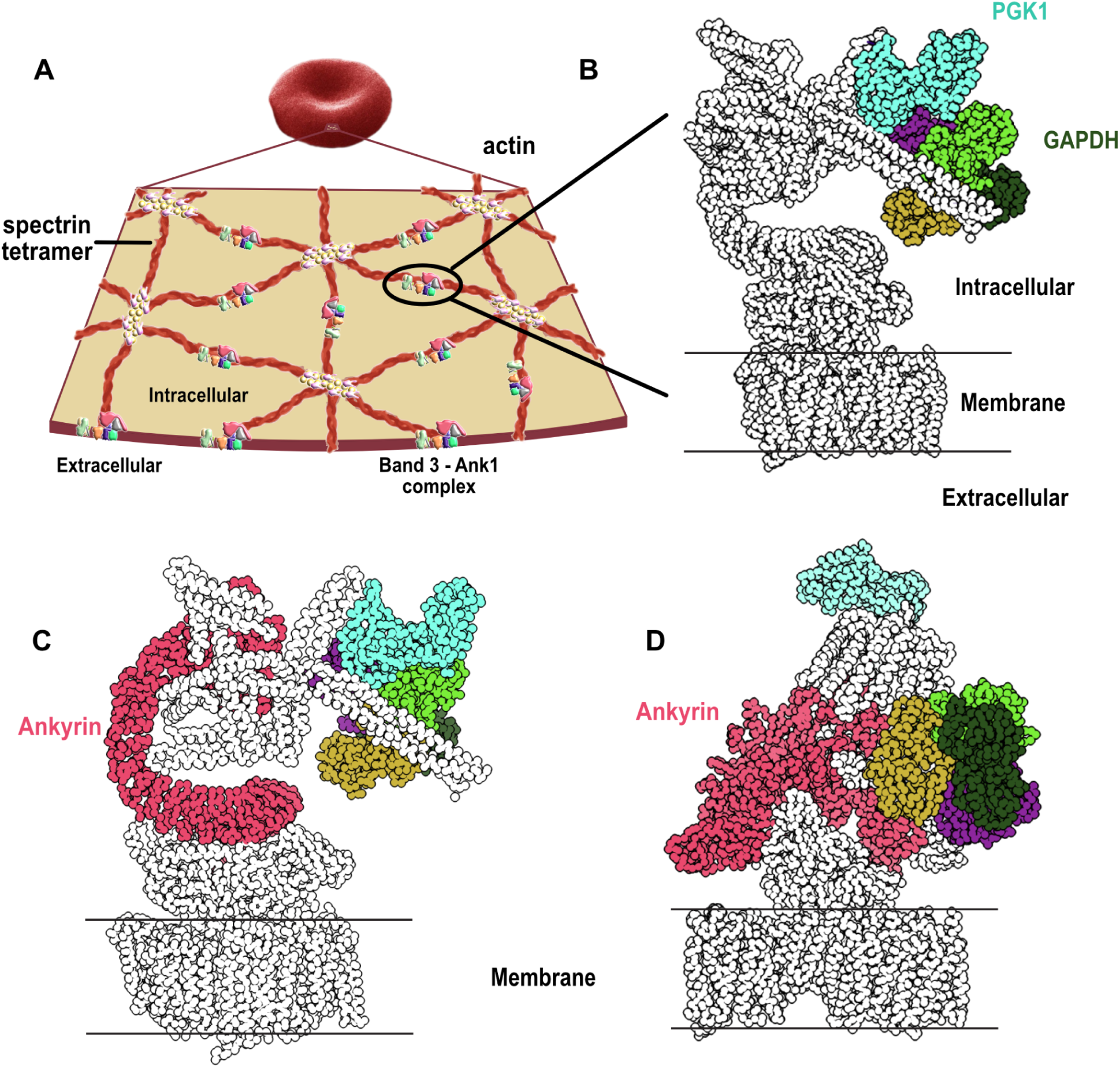
Reconstructions of band 3-Ank1-accessory protein complexes by integrative 3D modeling suggest Ank1 compression links the membrane to the cytoskeleton. (A) An overview of the cytoskeletal network supporting the membrane of RBC. A pseudohexagonal network of spectrin heterotetramer underlies the membrane and is anchored to the membrane by the band 3-Ank1 complex. The tetramer is attached to actin on the other end through the association of band 4.1, actin, spectrin and other proteins. An actin polymer can interact with 6 spectrin tetramers through band 4.1 (Lux, 2016) (adapted with permission from (Goodman, 2020)). (B) Glycolytic enzymes such as GAPDH and PGK1 are anchored to the band 3-Ank1 complex which can accommodate these enzymes while ANK1 adopts either open or closed conformations (see **Figure S13** for more details). (C) Ank1 in an open form in the band 3-Ank1 complex. Ank1, GAPDH, PGK1 are colored. (D) Roughly a third of observed intramolecular Ank1 crosslinks support it adopting a closed form *in situ* relative to the extended conformation observed for purified ankyrin (Wang et al., 2014), suggesting that Ank1 is capable of adopting either open or closed conformations. (see **Figure S11-12** for more details).

In addition to the membrane and cytoskeletal proteins, we found crosslinking evidence for glycolytic enzymes associating with the band 3-Ank1 complex. Based on our modeling, both the potential open and closed forms of ankyrin in the band 3 complex (**Figure 6B-D**) can accommodate glyceraldehyde-3-phosphate dehydrogenase (GAPDH) and phosphoglycerate kinase 1 (PGK1) binding without any violations of crosslinker distances. Thus, the positions of these enzymes with regards to the band 3-ankyrin complex do not seem to have an impact on the structural integrity of the complex as a whole. Previous experiments confirm the association of these two enzymes with the RBC membrane: GAPDH was found to be associated at the red cell membrane *via* immunofluorescence and binding assays (Rogalski et al., 1989), and PGK1 activity was detected in RBC white ghosts (Harris and Winzor, 1990; Ronquist and Ågren, 1966; Schrier, 1966). It is possible that localizing energy-producing enzymes to the RBC membrane where ion channels, transporters, and cytoskeletal network are located could be a way for RBCs to rapidly adjust the ratio of cell volume to surface area and change conformational states of cytoskeletal proteins such as ankyrin and spectrin in a timely manner. In turn, these rapid changes would facilitate dynamic morphological changes in response to the movement of RBCs through tissues and microvasculature.

Taken together, the band 3-ankyrin complex maintains structural integrity by linking the spectrin network to the membrane, while also serving as the hub for metabolic activities. These metabolic activities include energy metabolism (glycolytic enzymes) and ions and gas exchange (ammonium through Rh proteins and CO_2_, HCO_3_, Cl through band 3, and carbonic anhydrase (Sterling et al., 2001)). Such elaborate organization of channels, integral proteins, and enzymes makes RBCs a unique and minimalist system which can undergo rapid morphological changes while preventing cell lysis from occurring.

### Conclusions

We have used protein mass spectrometry, electron microscopy, chemical crosslinking, and 3D modeling to systematically determine the protein organization of the simplest primary cell in the human body, the erythrocyte. After defining a consensus set of canonical red blood cell proteins, we constructed a reference map of RBC protein interactions and multiprotein complexes, providing a detailed look at the molecular organization of RBC proteins. A 3D model of the band 3-Ank1 complex provides amino acid level detail of a critical protein complex that underlies one of RBCs most fascinating features: the ability of RBCs to undergo rapid dynamic morphological changes, a key aspect of controlling cellular form and activity given RBCs’ absence of gene expression regulation. The detailed molecular organization of proteins in the band 3-Ank1 complex clarifies the structure of a major membrane-cytoskeletal attachment site, supports the possibility of spring-like behavior at the membrane/cytoskeleton connection, and suggests aspects of RBC metabolism are spatially organized. In all, these data represent the first near-complete map of protein-protein interactions in any primary human cell type and thus provide an essential reference for these minimalistic cells, laying the groundwork for future full-cell molecular models.

## Materials and Methods

### Data and code availability statement

Raw and processed protein mass spectrometry data have been deposited in the MASSIVE/ProteomeXchange database in entry PXD030050, available at doi:10.25345/C5R00B. The 20S proteasome cryo-EM structure was deposited in the Electron Microscopy Data Bank as entry EMD-24822. Electron micrograph datasets have been deposited in EMPIAR as entry EMPIAR-10848. Supporting data files, including descriptions of biochemical fractionations, feature matrices, integrated 3D models, and training and test sets, have been deposited at Zenodo as accession doi:10.5281/zenodo.5768148.

### Native protein extraction

All blood cell types and plasma were purchased from Gulf Coast Regional Blood Center (Houston, TX). The most recently drawn blood was requested for each order of cells/plasma, and the orders were shipped on ice overnight to ensure cells’ freshness. Further steps (described below) were performed to improve cell type homogeneity. Typically, 1-4 mg of extract was 0.45μm filtered (Ultrafree-MC HV Durapore PVDF, Millipore) and fractionated by HPLC chromatography. Detergent and salt were minimized where possible to avoid perturbing protein assemblies. The protein concentration was determined by Bio-Rad Protein Assay or Bio-Rad DC Protein Assay (for samples containing detergent). For additional details on types of fractionations and sample-related information, see the Zenodo data repository.

### Red blood cells

According to the supplier, leukocytes were reduced via centrifugation of whole blood. However, we took additional steps to further eliminate contaminating plasma proteins and other cell types, namely leukocytes, reticulocytes, and platelets. Each blood unit was stored at 4 °C for 5 days after receipt in order to allow reticulocytes to mature into RBCs before extracting proteins (Urbina and Palomino, 2018). To further reduce leukocytes and platelets, RBCs were washed with PBS pH 7.4 (Gibco, Thermo Scientific, CA, USA) at 200 ×g for 15 minutes for 3 times. At the final wash, the 1 mm top layer of the cells (residual platelets, WBCs, etc) was removed. Cell samples and buffers were kept at 0−4 °C during all steps. Purified RBCs were lysed with 5 volumes of hypotonic lysis solution (5 mM Tris-HCl pH 7.4) containing EDTA-free protease inhibitor cocktail tablets (Sigma) and phosphatase inhibitor tablets (PhosSTOP EASY, Roche). White ghosts and hemolysate were separated from each other by centrifugation at 21 000 ×g for 40 min. The protein concentration of hemolysate was measured, and the samples were snap-frozen with liquid nitrogen until further use.

For ghosts preparation, excess hemoglobin (Hgb) was reduced by washing white ghosts ∼10 times with a hypotonic solution until the white ghosts appeared pale pink to almost white. Ghosts were then dissolved with the appropriate detergent (see Zenodo data repository for details), and the protein concentration was measured with a detergent-compatible Bio-Rad DC Protein Assay. For ghosts dissolved in Diisobutylene/Maleic Acid (DIBMA, Cube Biotech) (Oluwole et al., 2017), DIBMA was added to white ghosts to the final concentration 2.5% (50 mM HEPES pH 7.5), and the sample was rotated at 4 °C overnight (Hagn et al., 2018). The ghosts dissolved in 2.5% DIBMA were then treated as for other detergent-dissolved white ghosts. For hemolysate preparations, remnant white ghosts were removed by centrifugation at 21,000 ×g for 40 min at 4 °C, and the supernatant treated with Hemoglobind (Biotech Support Group) in order to bind and remove free Hgb. Protein concentrations of the Hgb-reduced samples were measured by Bradford colorimetric assay (Bio-Rad) prior to biochemical fractionation.

### iPSC-derived erythroblasts

Hematopoietic differentiation from iPSCs to hematopoietic stem and progenitor cells (HSPCs) was induced according to (Leung et al., 2018). To induce erythroid differentiation from HSPCs, day-15 HSPCs were specified using a 2-step suspension culture system consisting of Serum-Free Expansion Medium II, 2 mM of L-glutamine, and 100 mg/mL of primocin at 37°C in normoxic, 5% carbon dioxide conditions. Between days 15 and 20, this base was supplemented with 100 ng/mL of human stem cell factor, 40 ng/mL of IGF1, 5×10^-7^ M of dexamethasone, and 0.5 U/mL of hEPO and between days 20 and 25 with 4 U/mL of hEPO. Protein was extracted from day 25 differentiated cells, at which stage the differentiation cultures consisted of polychromatic and orthochromatic erythroblasts (Vanuytsel et al., 2018).

### Platelets

Prostaglandin E1 (PGE1), the aggregation inhibitor, was added to platelet rich plasma (PRP) to the final concentration of 1 mM in order to stop the activation of platelets. To remove contaminating red and white blood cells, contaminating RBCs and WBCs were pelleted at 100 x g for 10 mins at 4 °C with no brake applied. The supernatant (PRP) was then washed two times with 1 volume platelet-wash buffer : 1 volume PRP by centrifugation at 400 x g for 10 min at 4 C with no brake. The wash buffer contains 1 μM PGE1, 10 mM sodium citrate, 150 mM NaCl, 1 mM EDTA, 1% (w/v) glucose pH 7.4. Native protein complexes from platelets were obtained by incubating the washed pallet with Pierce IP lysis buffer (25 mM Tris•HCl pH 7.4, 150 mM NaCl, 1% NP-40, 1 mM EDTA, 5% glycerol) (Thermo Scientific) containing protease and phosphatase inhibitors at 4 °C for 5 minutes. The lysate was clarified at 14,000 x g for 10 mins.

### Plasma

Plasma separated from whole blood by centrifugation was purchased and shipped in anticoagulant CPD. The received plasma still contained other blood cell types, so these cells were removed through centrifugation. Plasma was centrifuged at 5,000 x g for 15 mins at 4°C, and only the top part of supernatant was collected. The supernatant was treated with Albuviod (Biotech Support Group) to reduce the presence of serum albumin (HSA). The HSA reduced supernatant was then concentrated in a 3 kD MWCO Amicon Ultra filter unit (MilliporeSigma, Burlington, MA).

### White blood cells

In order to broadly assess WBC proteins, we analyzed a buffy coat, consisting of all WBCs (lymphocytes and granulocytes), as well as some RBCs and platelets. WBCs were separated from the other cell types using Histopaque 1077 (Sigma) according to the manufacturer’s protocol. Briefly, the buffy coat was diluted with PBS Gibco at a 1:1 ratio. Buffy coat and Histopaque were held at room temperature to ensure proper cell separation. PGE1 was added to the diluted buffy coat to a final concentration of 1 μM to prevent platelet activation and cell aggregation. WBCs were first purified using Histopaque 1077 where the “ring” of lymphocytes, the layer between plasma and histopaque after centrifugation, and the white layer containing granulocytes on top of the RBC layer at the bottom were kept for further purification steps. Contaminating RBCs were lysed by addition of hypotonic solution, and the mixture of lymphocytes and granulocytes were pelleted at 1,000 g for 10 minutes. Native protein extract of WBCs was prepared by adding Pierce IP lysis buffer, and the extract was clarified by centrifuging at 14,000 x g for 10 mins, prior to biochemical fractionation.

### HPLC chromatography

Hgb-reduced hemolysate and detergent-dissolved ghosts were fractionated on a Dionex UltiMate3000 HPLC system consisting of an RS pump, Diode Array Detector, PCM-3000 pH and Conductivity Monitor, Automated Fraction Collector (Thermo Scientific, CA, USA) and a Rheodyne MXII Valve (IDEX Health & Science LLC, Rohnert Park, CA) using biocompatible PEEK tubing and various columns. The columns we used were size exclusion chromatography, ion exchange separations (mixed bed or triple-phase WAXWAXCAT), or hydrophobic interaction chromatography. The sample loaded was 1-4 mg protein as measured by the BioRad Protein Assay (hemolysate) or DC Protein Assay (detergent-dissolved ghosts) as appropriate to the sample buffer. Fractions were collected into 96-deep well plates.

Size exclusion: BioSep-SEC-s4000 600 x 7.8 mm ID, particle diameter 5 μm, pore diameter 500 Å (Phenomenex, Torrance, CA) or higher Mw BioBasic AX HPLC Columns, 600 x 7.8 mm ID 5μm particle size, 300Å pore size were used. Unless otherwise specified the sample was 200 μl, low rate 0.5 ml/min, with fraction collection every 45 seconds, and mobile phase was PBS pH 7.4 (Gibco). For column calibration, molecular weight standards (Sigma -Aldrich, MWGF1000, 2-5 ug each of carbonic anhydrase (C7025), β-amylase (A8781), and bovine serum albumin (A8531)) were fractionated to ensure that the column can separate proteins based on their sizes correctly before the hemolysate/ghosts samples were fractionated.

Mixed bed ion exchange: Poly CATWAX A (PolyLC Mixed-Bed WAX-WCX) 200 x 4.6 mm ID, Particle diameter 5 μm, pore diameter 100 Å (PolyLC Inc., Columbia, MD). The bed contains the cation-exchange (PolyCAT A) and anion-exchange (PolyWAX LP) materials in equal amounts. A 200-250 μl sample was loaded at ≤ 40 mM NaCl, and eluted with a 1-hour salt gradient at 0.5 ml/min with collections of 0.5 ml fractions. Gradient elution was performed with Buffer A (10 mM Tris-HCl pH 7.5, 5% glycerol, 0.01% NaN3), and 0-70% Buffer B (1.5 M NaCl in Buffer A).

Triple phase ion exchange (WWC): Three columns, each 200 x 4.6 mm ID, particle diameter 5 μm, pore diameter 100 Å, were connected in series in the following order: two PolyWAX LP columns followed by a single PolyCAT A (PolyLC, Inc, Columbia, MD). Loading, buffers, and fraction collection were as for mixed bed ion exchange above with slight modifications in low rate and elution from the methods of (Havugimana et al., 2012). The low rate was either 0.25 ml/min with a 120 min gradient from 5-100 %B. For the separation of nuclear extracts, the gradient was modified to a 115-minute multiphasic elution from 5-100% Buffer B.

Hydrophobic interaction: The ProPac HIC-10 hydrophobic interaction chromatography column (Thermo Scientific) had the following characteristics: 4.6 x 250 mm, Amide/Ethyl (phase), 5 µm particle size, and 300 Å pore size. A 200-250 μl sample was loaded at ∼250 mM (NH_4_)_2_ SO_4_, and eluted with a 100-min low salt gradient at 0.5 ml/min with collections of 0.5 ml fractions. Gradient elution was performed with Buffer A (2 M (NH_4_)_2_ SO_4_ in 0.1M NaH_2_PO_4_ pH 7.0), and 0-70% Buffer B (0.1 M NaH_2_PO_4_ pH 7.0).

### Mass spectrometry sample preparation

Samples were prepared for mass spectrometry in 96-well plate format using ultrafiltration and in-solution digestion protocols (Wan et al., 2015). Plates were sealed with transparent film during incubation steps.

Ultrafiltration was performed with an AcroPrep Advance 96-filter plate, 3kD MWCO (Pall) using a vacuum manifold (QIAvac 96 or Multiwell, Qiagen) at -0.75 Bar. Before filtering samples, preservatives and remnant polymers that could interfere with MS experiment were removed from the plate by sequential filtration of 100 μl LC/MS quality water and 100μl trypsin digestion buffer (50 mM Tris-HCl, pH 8.0, 2 mM CaCl2). Samples were concentrated to 100 μl, diluted 2-fold with trypsin digestion buffer, and concentrated again to a final volume of 50-100 μl before being transferred back to a 96-deep well plate. 50 μl 2,2,2,-trifluoroethanol (TFE) was added and samples were reduced with TCEP (Bond-Breaker, Thermo) at a final concentration of 5 mM for 30 min. 37°C. Iodoacetamide was added to 15 mM and plates were incubated in the dark 30 min at room temperature. Alkylation was quenched by the addition of DTT to 7.5 mM. TFE was diluted to <5% by the addition of trypsin digestion buffer. 1 μg of trypsin was added to each fraction and the sealed plate was incubated 37°C overnight. Digestion was stopped by adding formic acid to the final concentration of 0.1%, and peptides were desalted using a 5-7 μl C18 Filter Plate (Glygen Corp) with a vacuum manifold, dried, and resuspended for mass spectrometry in 3% acetonitrile, 0.1% formic acid.

### Chemical cross-linking

Extract (∼2-3 mg hemolysate and detergent-dissolved white ghosts) was fractionated by size exclusion as written above. DSSO (Thermo Scientific) was dissolved immediately before use in dry dimethylformamide or DMSO (stored under nitrogen) at a concentration of 50 mM and then further diluted in PBS for a working stock. Immediately after fractionation, working stock DSSO was added to fractions to a final concentration of 1 mM, and samples were incubated 1 hour at room temperature. Cross-linking was quenched by the addition of 1 M Tris pH 8.0 to a concentration of 24 mM. Cross-linked fractions were immediately frozen and stored at -80°C. Fractions were prepared for mass spectrometry using the ultrafiltration and in-solution digestion methods. Peptides were desalted before being analyzed by mass spectrometry.

### Mass spectrometry data acquisition and processing

#### Acquisition

Mass spectra were acquired using one of two Thermo mass spectrometers: Orbitrap Fusion or Orbitrap Fusion Lumos. In all cases, peptides were separated using reverse phase chromatography on a Dionex Ultimate 3000 RSLCnano UHPLC system (Thermo Scientific) with a C18 trap to Acclaim C18 PepMap RSLC column (Dionex; Thermo Scientific) configuration. Peptides were eluted using a 3-45% gradient over 60 min. (for Fusion and Lumos) and directly injected into the mass spectrometer using nano-electrospray for data-dependent tandem mass spectrometry.

Mass spectrometry data was acquired for each instrument as follows:

#### Orbitrap Fusion

top speed CID with full precursor ion scans (MS1) collected at 120,000 m/z resolution and a cycle time of 3 sec. Monoisotopic precursor selection and charge-state screening were enabled, with ions of charge > + 1 selected for collision-induced dissociation (CID). Dynamic exclusion was active with 60 s exclusion for ions selected once within a 60 s window. For some experiments, a similar top speed method was used with dynamic exclusion of 30 s for ions selected once within a 30 s window and high energy-induced dissociation (HCD) collision energy 31% stepped +/-4%. All MS2 scans were centroid and done in rapid mode.

#### Orbitrap Lumos

top speed HCD with full precursor ion scans (MS1) collected at 120,000 m/z resolution. Monoisotopic precursor selection and charge-state screening were enabled using Advanced Peak Determination (APD), with ions of charge > + 1 selected for high energy-induced dissociation (HCD) with collision energy 30% stepped +/- 3%. Dynamic exclusion was active with 20 s exclusion for ions selected twice within a 20s window. All MS2 scans were centroid and done in rapid mode.

For identification of DSSO cross-linked peptides, peptides were resolved using a reverse phase nano low chromatography system with a 115 min 3-42% acetonitrile gradient in 0.1% formic acid. The top speed method collected full precursor ion scans (MS1) in the Orbitrap at 120,000 m/z resolution for peptides of charge 4-8 and with dynamic exclusion of 60 sec after selecting once, and a cycle time of 5 sec. CID dissociation (25% energy 10 msec) of the cross-linker was followed by MS2 scans collected in the orbitrap at 30,000 m/z resolution for charge states 2-6 using an isolation window of 1.6. Peptide pairs with a targeted mass difference of 31.9721 were selected for HCD (30% energy) and collection of rapid scan rate centroid MS3 spectra in the Ion Trap.

### Computational analyses of peptide mass spectra

The human reference proteome was downloaded from Uniprot.org (UniProt Consortium, 2019) in August 2018. Mass spectral peptide matching was performed with MSGF+, X!Tandem, and Comet-2013020, each run with 10 ppm precursor tolerance, and allowing for fixed cysteine carbamidomethylation (+57.021464) and optional methionine oxidation (+15.9949). Peptide search results were integrated with MSBlender (Kwon et al., 2011), https://github.com/marcottelab/msblender, https://github.com/marcottelab/run_msblender). For DSSO cross-linked experiments, inter-protein cross-links were identified using the XlinkX (Klykov et al., 2018) node in ProteomeDiscover 2.2 (ThermoScientific). FDR for all analyses was kept at 1%.

### RBC proteome

#### Identification of RBC proteome

We constructed a supervised classifier, trained on proteomics and RNA-seq data of different blood cell types (WBCs, platelets, plasma proteins, RBC precursor cells), in order to comprehensively identify the proteome of RBCs. MS data of different blood cell types were retrieved from the PRIDE database; several of the datasets were reanalyzed using either ProteomeDiscover 2.2 or MSBlender as noted in the Zenodo data repository. A classifier feature matrix was assembled wherein rows were proteins observed in MS and RNA-seq experiments, corresponding to columns, and matrix elements were peptide-spectrum matches (PSM) and reads per kilobase (RPKM), as appropriate. As gold standard RBC proteins, we used the set of 859 proteins that all three previous MS studies on RBCs identified (Bryk et al., 2017, Goodman et al., 2007 and Lange et al., 2014) as a positive set (*i.e*., most likely to be true RBC proteins). As a negative gold standard (*i.e*., most likely *not* RBC proteins), we selected all proteins found in the whole human proteome curated in Uniprot (20,350 reviewed proteins) but not found in any of the three prior RBC studies (3,520 proteins found in the three studies). Proteins found in at least one but not all 3 prior RBC studies were considered as unknowns to be classified. We divided the gold standard proteins into train and test sets (80:20, train:test), then used TPOT, an autoML wrapper of scikit-learn machine learning functions (Olson et al., 2016), to perform all training steps, including identifying the best classifier and hyperparameters based on 5-fold cross-validation. Application of that model to the entire set of proteins gave confidence scores for each protein to be an RBC protein. Precision and recall were calculated from training (687 positive, 13,665 negative) and test (172 positive, 3,417 negative) set proteins for **Figure 1a** and **1b**, respectively. The false discovery rate was calculated from the full set of 859 positive and 17,082 negative proteins.

### Defining RBC complexes based on the CF-MS datasets

#### Assembly of features for scoring putative protein interactions

For each experiment, we assembled an elution matrix of all identified proteins by fractions with PSMs normalized to 1 within each protein. In addition, we concatenated elution matrices of all 1,944 fractionation experiments as one additional matrix . Therefore, the total number of matrices used to calculate pairwise scores was 30. Cross-linked data of hemolysate and ghosts were not used in training. We trained only proteins within the determined RBC proteome (see above) at 5% FDR. Next, we calculated a series of all-by-all pairwise scores between proteins for individual matrices and the concatenated matrix. The scores/features are as follows: (1) Pearson’s r, (2) Spearman’s rho, (3) Euclidean distance, (4) Bray-Curtis similarity, (5) stationary cross correlation, (6) covariance, and (7) hypergeometric score for the co-occurrence of proteins in fractions with repeated sampling of fractions (Drew et al., 2017). All features/scores were calculated with added Poisson noise. Euclidean distance and Bray-Curtis similarity scores were inverted and normalized to a max score of 1.

For each individual matrix from each fractionation, features 1-6 were calculated. Additionally, the average of each type of score/feature was calculated from all individual matrices excluding the concatenated matrix. For the concatenated matrix of all fractionations, features 1-7 were calculated. Features were joined to create a final matrix composed of 4,131,128 rows and 193 features. Missing values were filled with zeros.

#### Construction of the gold standard protein complex training and test sets

We used known human protein complexes from the CORUM database (Giurgiu et al., 2019)) as a gold standard set of positive stable protein-protein interactions. 204 known complexes were divided into positive training and test complexes according to the scheme from (Drew et al., 2017) and complexes with over 30 members were removed. 25,797 negative training and test interactions were drawn from feature matrix rows, removing any interactions present in the positive gold standard set. 461 positive training interactions and 299 positive test interactions were present in the feature matrix.

#### Identification of interacting proteins by supervised machine learning

We again utilized TPOT to train our machine learning model to find pairwise interactions among members of protein complexes. We discovered optimal hyperparameters for an ExtraTree with 5-fold cross-validation, with an area under the precision-recall curve of 0.45. We then trained an ExtraTree with TPOT discovered hyperparameters, and the resulting model was applied to the entire feature matrix to give a CF-MS score to each pair of proteins, with higher scores corresponding to higher confidence in the proteins interacting (specifically, being subunits in the same multi-protein assembly). Precision, recall, and false discovery rates were calculated from 299 positive and 11,408 negative test set interactions.

### Negative stain electron microscopy

4 μL of hemolysate was applied to a glow-discharged 400-mesh continuous carbon grid. After allowing the sample to adsorb for 1 min, the sample was negatively stained with five consecutive droplets of 2% (w/v) uranyl acetate solution, blotted to remove residual stain, and air-dried in a fume hood. Grids were imaged using an FEI Talos TEM (Thermo Scientific) equipped with a Ceta 16M detector. Micrographs were collected manually using TIA v4.14 software at a nominal magnification of x73,000, corresponding to a pixel size of 2.05 Å/pixel. CTF estimation, particle picking, and 2D class averaging were performed using both RELION v3 (Zivanov et al., 2018) and cryoSPARC v2.12.4 (Punjani et al., 2017). Three negative stain datasets were collected. The first dataset collected contained ∼220 micrographs of pooled HPLC size exclusion fractions 1-9. ∼2,500 particles were manually picked and processed in cryoSPARC to produce the TPP2 structure in **Figure 3B-C**. Two datasets were collected of hemolysate after being passed through a 100 kDa filter, one as prepared and the other at 1:100 dilution. For the diluted sample, ∼400 micrographs were collected and ∼42,500 particles were picked using Topaz (Bepler et al., 2019). The resulting particles were used to generate the PRDX2 structure in **Figure 3B-C** and the 2D class averages in **Figure S3B**. For the non-dilute sample, ∼230 micrographs were collected and ∼1,500 proteasome particles were manually picked followed by classification in RELION **Figure S3C**.

### Cryo-EM grid preparation and data collection

C-flat holey carbon grids (CF-1.2/1.3, Protochips Inc.) were glow-discharged for 1 min using a Solarus 950 plasma cleaner (Gatan). 2 μL of 0.2 mg/mL graphene oxide (Sigma-Aldrich) was placed onto the grids for 1 min followed by one wash with water. 3 μL of pooled and concentrated hemolysate from HPLC size exclusion fractions 10-20 was placed onto the grid, blotted for 3.5 sec with a blotting force of 0, and rapidly plunged into liquid ethane using an FEI Vitrobot MarkIV operated at 4 °C and 100% humidity. Data was acquired using an FEI Titan Krios TEM (Sauer Structural Biology Laboratory, University of Texas at Austin) operated at 300 keV with a nominal magnification of ×22,500 (1.045 Å/pixel) and defocus ranging from –1.09 to –2.5 μm. Dose-fractionated movies were collected using 20 frames (0.15 sec/frame) over a total of 3 sec with a dose rate of ∼2.13 e^-^/Å^2^/sec and a total exposure of 42.58 e^-^/Å^2^. A total of 6,606 micrographs were automatically recorded on a K3 detector (Gatan) operated in counting mode using Leginon (Suloway et al., 2005). A full description of the cryo-EM data collection parameters can be found in **Table S1**.

### Cryo-EM data processing

Motion correction, CTF-estimation and particle picking were performed in Warpv1.0.7 (Tegunov and Cramer, 2019). ∼1,000,000 extracted particles were imported into cryoSPARC v2.12.4 for 2D classification, 3D classification and non-uniform 3D refinement (**Figure S2)**. ∼60,000 particles were used in the final 20S proteasome reconstruction. The nominal resolution of the map using the gold-standard Fourier Shell Correlation (FSC) at 0.143 is 3.35 Å **(Figure S2)**. A previously solved X-ray crystal structure of the human 20S proteasome, PDB 6RGQ (Rêgo and Fonseca, 2019), was aligned by cross-correlation in UCSF Chimera (Pettersen et al., 2004) and used as an initial model for refinement. The model was then refined through two rounds of molecular dynamics flexible fitting and real space refinement using Namdinator (Kidmose et al., 2019), followed by further refinement in Phenix (Adams et al., 2002).

### Integrative 3D modeling

Integrative modeling of the red blood cell membrane complexes consisted of four main stages: 1) gathering data, 2) domain representation and configuring of spatial restraints, 3) system sampling and scoring of restraints, and 4) model validation, as previously described in integrative modeling work (Gutierrez et al., 2020; Kim et al., 2018; Russel et al., 2012). The python interface of the Integrative Modeling Platform (IMP) was used to model the complexes (Saltzberg et al., 2019) and all associated data, scripts, and outputs can be found at Zenodo data deposit. The following sections describe the method for modeling the ankyrin complex in the open-spring conformation, the ankyrin complex in the closed-spring conformation, the ankyrin complex in the open-spring conformation with metabolic enzymes bound, the ankyrin complex in the closed-spring conformation with metabolic enzymes bound, and the band 4.1 with spectrin subcomplex.

#### Data used for modeling

Protein structure representations were constructed from known X-ray crystal structures, modeled using I-TASSER (Yang et al., 2015), or modeled domains from the Swiss-Prot database. **Table S3** indicates the source of each of the protein’s structural models. A total of 156 intra- and intermolecular DSSO crosslinks were used to model the ankyrin complex. An additional 21 DSSO crosslinks were used to incorporate the GAPDH and PGK1 enzymes into the model. The subcomplex of spectrin and band 4.1 was modeled using 41 intra- and intermolecular DSSO crosslinks. Transmembrane regions of proteins were determined from Uniprot (The UniProt Consortium, 2019) annotations.

#### Domain representation and configuring spatial restraints

Protein subunits were represented as rigid bodies, chains of rigid bodies, or beads (**Table S3**). Band 3 was represented using rigid bodies for the transmembrane region and cytoplasmic domain of the protein. These domains were connected with a chain of flexible beads. The transmembrane region of the band 3 dimer (Arakawa et al., 2015) was represented as a single rigid body. Glut1 and band 4.2 were represented as a single rigid body, due to their high C-score (**Table S3**) from I-TASSER and agreement with intramolecular crosslinks. The transmembrane regions of the GYPA dimer have a known NMR structure and are represented as a single rigid body. The cytoplasmic and extracellular domains of the GYPA dimer were represented using a flexible chain of beads. RhAG and the two RhCE/D proteins were superimposed on the x-ray crystal structure (PDB 3HD6) and treated as a single rigid body (Gruswitz et al., 2010); these proteins had high C-scores and satisfied intermolecular crosslinks between each other. ANK1 was represented as a chain of rigid bodies (both modeled and X-ray crystal structure) with the end represented as flexible beads. Residues 265 to 790 of SPTA1 were used in the model and were represented as a chain of rigid bodies, with each rigid body beginning/ending at the Uniprot annotated spectrin domains. Residues 1,585 to 2,006 of SPTB were represented similarly to SPTA1 with each annotated spectrin domain being a rigid body connected in a chain. Both GAPDH and PGK1 had available X-ray crystal structures that satisfied all intramolecular crosslinks and were represented as rigid bodies. For the band 4.1-spectrin complex, band 4.1 was represented as a single rigid body, while residues 1,929 to 2,386 of SPTA1 was represented as a chain of rigid bodies by spectrin domain and residues 46 to 741 of SPTB was represented as a chain of rigid bodies by spectrin domain. The flexible chains of beads were coarse-grained to 10 residues per bead.

The DSSO crosslinkers were modeled using a length of 21 Å. The excluded volume restraint was applied to the 10 residue beads, preventing volumes from occupying the same space. The sequence connectivity restraint was applied between beads. Based on Uniprot annotations, segments of proteins were labeled as either inside the membrane (transmembrane), above the membrane (extracellular), or below the membrane (cytoplasmic) scored using a sigmoid potential. All of these restraints were incorporated into the scoring framework for the model.

#### System sampling and scoring of restraints

The 38 rigid bodies were first randomized in an initial configuration, followed by a steepest descent minimization based on connectivity to ensure that neighboring residues are close together and Monte Carlo sampling. This sampling produced 600,000 models from 20 independent runs (unique starting positions). (Note that detailed explanations of assessing sampling exhaustiveness can be found in previous literature (Viswanath et al., 2017)). The ensembles were clustered based on their scoring parameters, including the clustering precision, which describes the variability between models in the cluster (**Table S2**). The best scoring cluster was selected to proceed (**Table S2**).

#### Model validation

The model cluster was first assessed against the input data. A crosslink is considered to be satisfied if any model in the cluster has the crosslink distance less than 40 Å. The crosslinks that are unsatisfied are believed to represent another conformational state of the complex. The convergence is determined through assessing the exhaustiveness of the sampling (Viswanath et al., 2017). This protocol tests convergence of the model score, whether the model scores were drawn from the same parent distribution, whether the structural clusters include models from each sample proportional to their size (chi-squared and sampling precision), and structural similarity between the model samples (**Table S2** CCC between two sample densities).

## ACKNOWLEDGEMENTS

The authors gratefully acknowledge Dr. Zhongwu Zhou for aid in cryo-EM sample preparation and data collection and Roden Luo for critical reading. Research was funded by grants from the National Institute of General Medical Sciences, Grant/Award Numbers: GM122480, R35GM138348; Cancer Prevention and Research Institute of Texas, Grant/Award Number: RR160088; National Science Foundation, Grant/Award Number: 2019238253; National Institutes of Health, Grant/Award Numbers: HD085901, DK110520; Army Research Office, Grant/Award Number: W911NF-19-1-0021; Welch Foundation, Grant/Award Numbers: W911NF-15-1-0120, F-1515, F-1938.

## AUTHOR CONTRIBUTIONS

Conceptualization and Methodology, WSL, EMM; Software, WSL, CLM, EJV, CDM, KD; Investigation, WSL, CLM, EJV, PCH, OP; Analysis - WSL, CLM, EJV Writing – Original Draft, WSL, EMM, CLM, EJV; Writing – Review & Editing, WSL, EMM, CLM, AE, KV, DWT; Funding Acquisition, EMM; Erythroblast analyses, KV, GM, PCH, AE; Resources, DWT, EMM; Supervision, EMM

---

**Figure S1.**
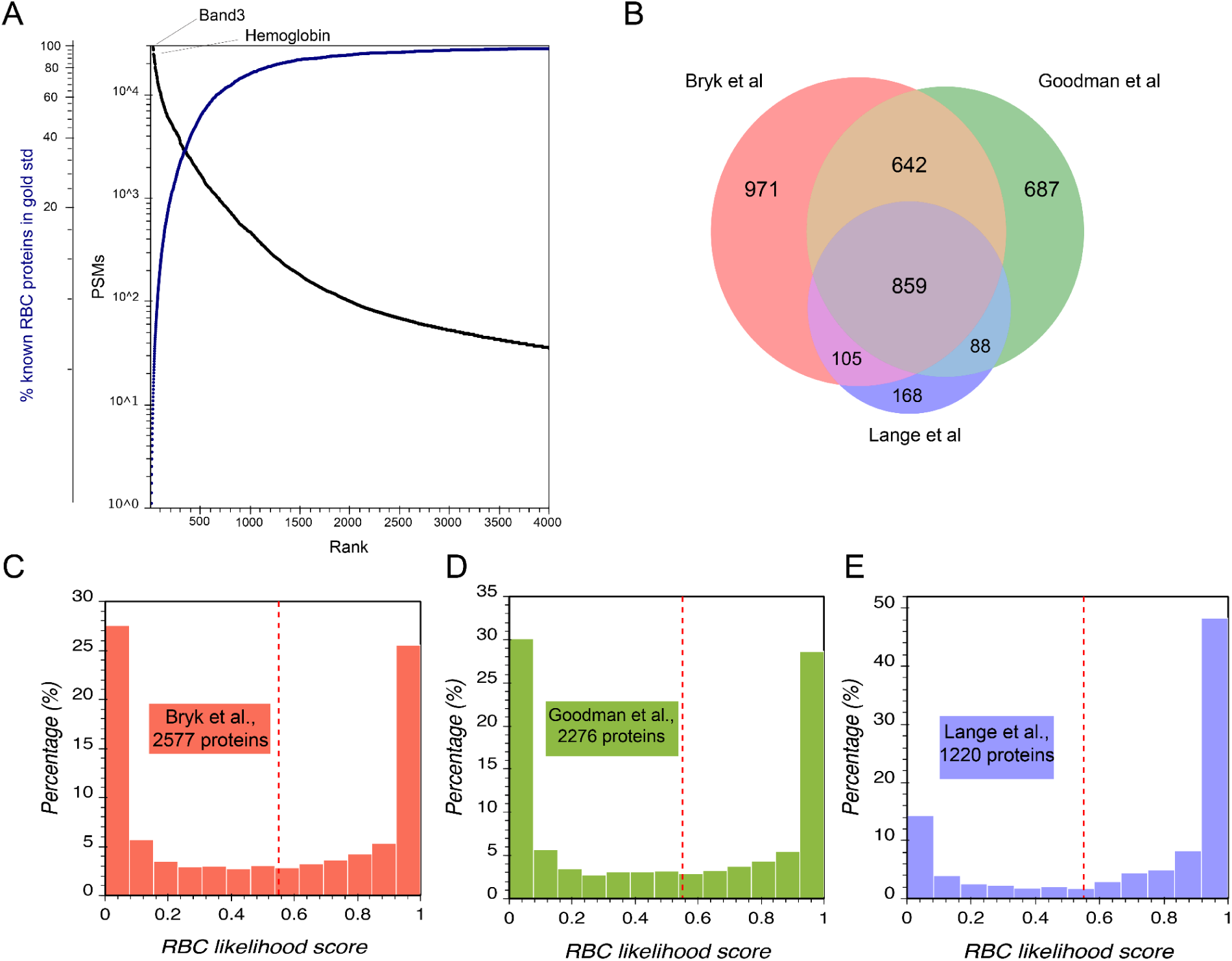
Coverage and Validation of RBC proteome. **(A)** RBC proteins were ranked based on abundance (black plot). While hemoglobin is the most abundant protein in RBCs, band 3 is the most abundant in our experiment because of hemoglobin depletion steps in our protocol. Blue plot shows >90% of the RBC proteins in the gold standard set of 859 proteins were detected within the most 2,000 abundant proteins in our experiment, showing a good coverage of RBC proteome. **(B-E)** A Venn diagram shows the overlap and differences of proteins detected in the three prior proteomic studies (Bryk and Wiśniewski, 2017; Goodman et al., 2007; Lange et al., 2014). Proteins in the intersection were considered well-supported RBC training examples; human proteins not observed by all 3 prior studies were considered non-RBC proteins. Proteins observed by only 1 or 2 prior studies were considered candidate RBC proteins to be scored by the classifier. The remaining panels plot the percentage of proteins in each of the 3 prior studies as a function of the confidence scores assigned by the classifier. Vertical red dashed lines indicate the 1% FDR threshold.

**Figure S2.**
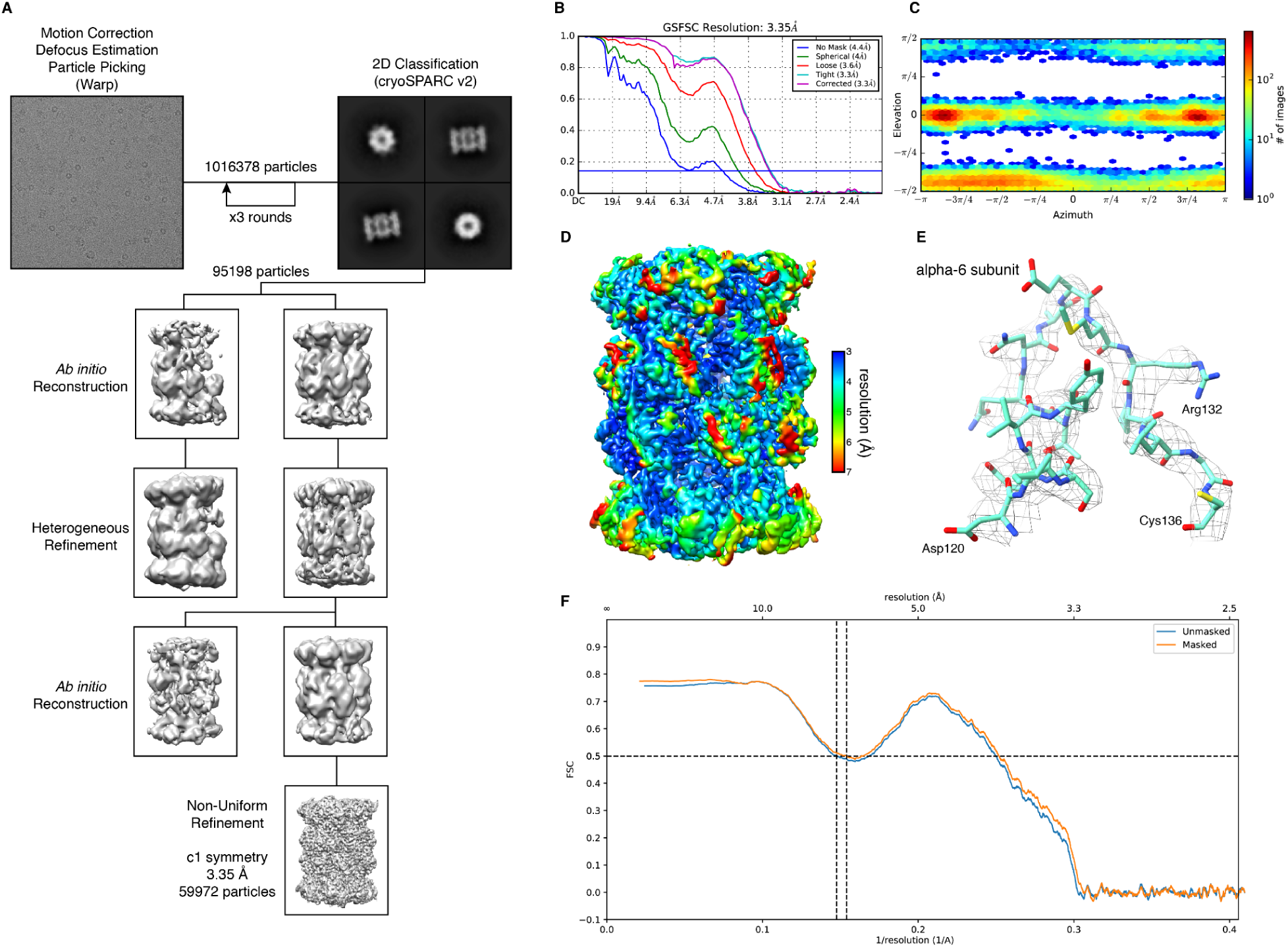
Cryo-EM data processing workflow and structure validation for 20S proteasome. **(A)** Cryo-EM data processing workflow for 20S proteasome. **(B)** Fourier Shell Correlation (FSC) curves for the 20S proteasome based on the gold-standard between two independent half maps. **(C)** Euler angle distribution plot for the 20S proteasome. **(D)** Local resolution map of the 20S proteasome reconstruction. **(E)** Region from the alpha-6 subunit of the 20S proteasome reconstruction and model showing a more highly resolved portion of the map. **(F)** Map-to-model FSC.

**Figure S3.**
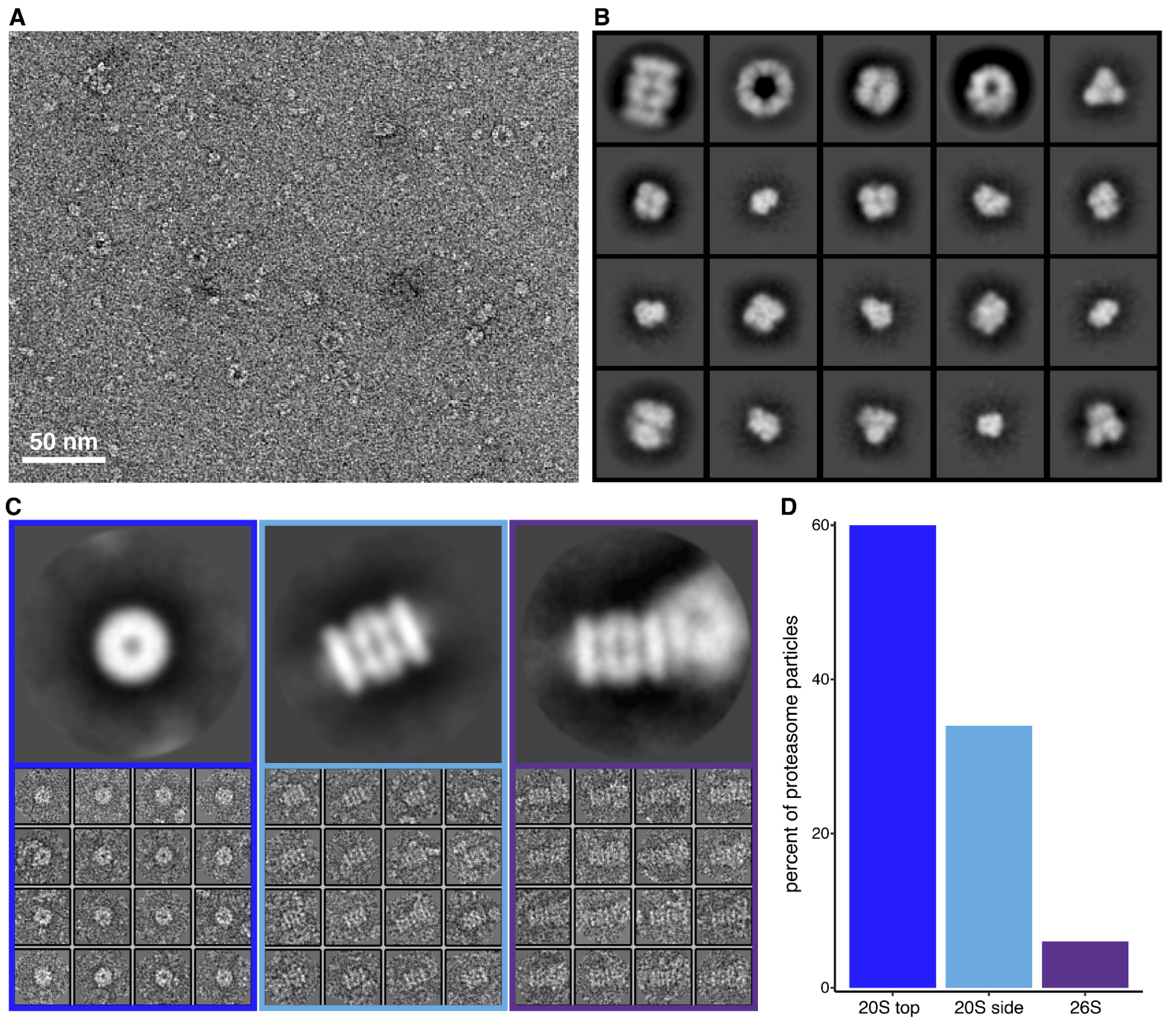
Assessment of proteasomes from negative stain electron microscopy of RBC hemolysate shows a majority in the 20S form. **(A)** Representative micrograph of hemolysate after being passed through a 100 kDa filter. **(B)** Representative reference-free 2D class averages from filtered hemolysate showing macromolecular assemblies of distinct sizes and shapes. Box length is 254 Å. **(C)** Reference-free 2D class averages and aligned raw particles for 20S proteasome (top view), 20S proteasome (side view) and 26S proteasome (single-capped) from left to right. Box length corresponds to 459 Å. **(D)** Distribution of observed proteasome states from negative stain EM of hemolysate. The total number of proteasome particles classified was 1,510.

**Figure S4.**
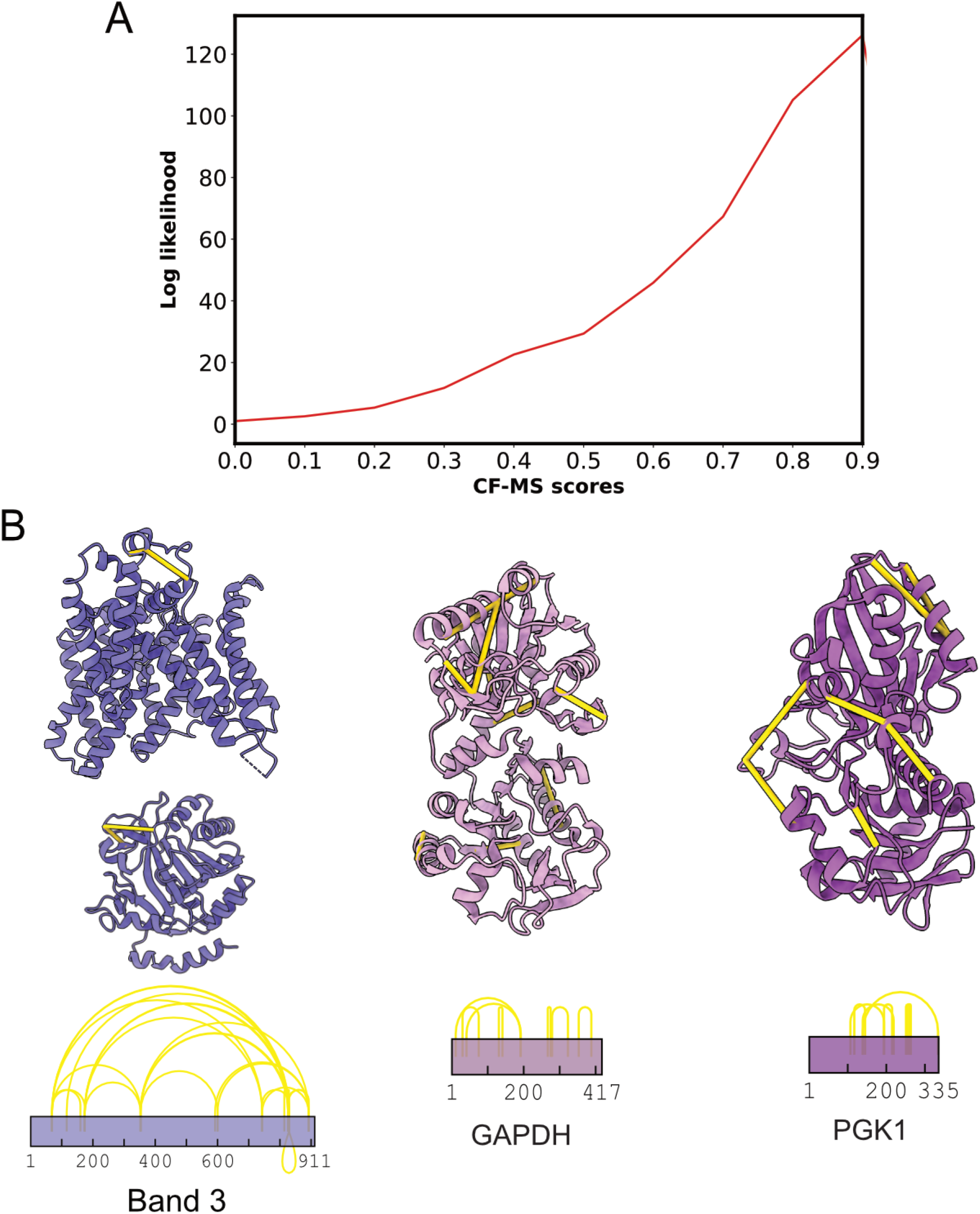
3D models of individual membrane/cytoskeletal proteins prior to integration into larger complexes. Yellow lines in the protein structures represent intramolecular crosslinks detected in our crosslinking experiments. The purple lines above rectangular boxes show the amino acid residue crosslinks detected. Intramolecular crosslinks of individual proteins agreed with the existing structures of these proteins.

**Figure S5.**
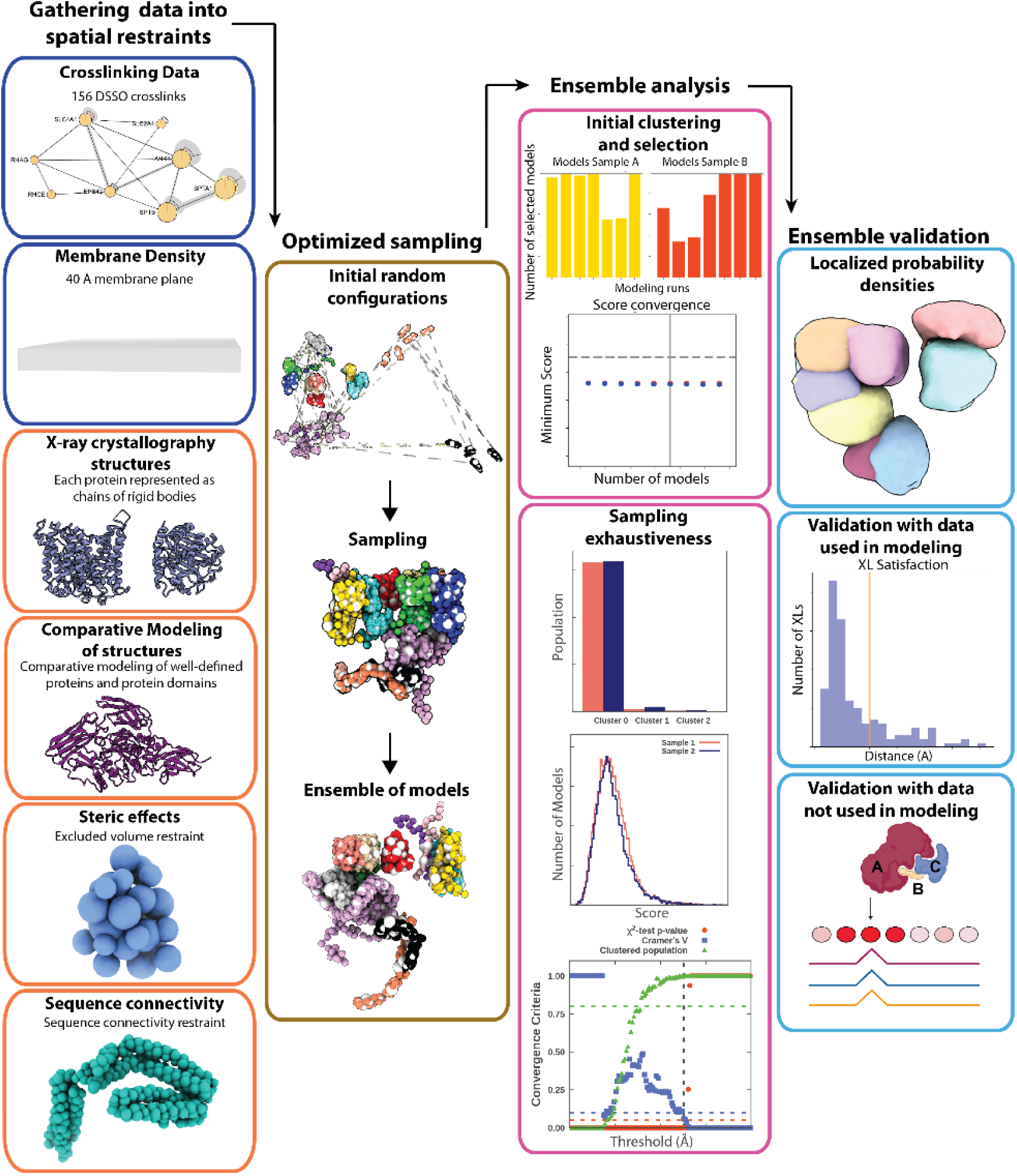
Integrative Modeling Platform (IMP) workflow. Data in the blue box are converted into spatial restraints. The orange boxes show x-ray crystal structures, statistical inferences, and physical properties. The gold box displays the sampling and scoring of the models.The models are analyzed via an initial clustering to select a high scoring group of models from various simulation runs followed by sampling exhaustiveness (pink boxes). The light blue boxes show the ensemble validation against the data used in the modeling, the probability density for each subunit, and a comparison against the known structure.

**Figure S6.**
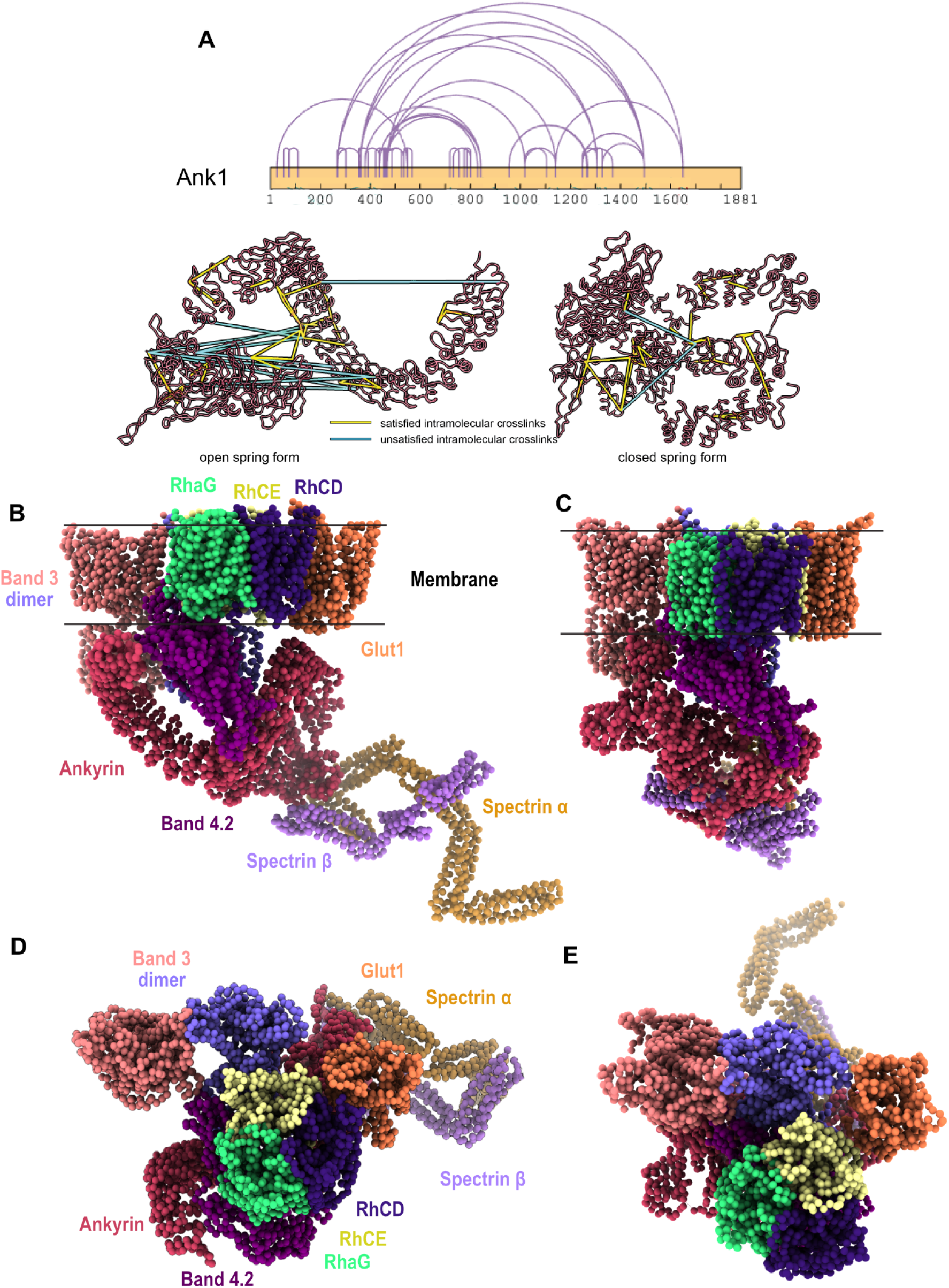
Correlated cross-linker violations suggest ankyrin may adopt a highly compressed structure within the band 3-Ank1 complex. **(A)** 30% of crosslinks exceed the inter-amino acid distances expected for the DSSO crosslinker (30 Å) when mapped onto the ANK1 structure in its open conformation (as observed crystallographically by (Wang et al., 2014), but these crosslinks can be satisfied by a model of ANK1 in a closed conformation, suggesting that ANK1 in the band 3-Ank1 complex may adopt both open and closed conformations. **(B&C)** Integrative 3D model of band 3-Ank1 complexes with Ank1 in open (left) and closed (right) conformations. Side and top views of the band 3-Ank1 complex with Ank1 in open conformation. **(D&E)** Side and top views of band 3-Ank1 complex with Ank1 in closed conformation.

**Table S1.** Cryo-EM data collection and reconstruction statistics.

|  |  |
| --- | --- |
| Protein | 20S Proteasome |
| EMDB | EMD-24822 |
| Microscope | FEI Titan Krios |
| Voltage (kV) | 300 |
| Detector | Gatan K3 |
| Magnification (nominal) | 22,500 |
| Pixel size (Å/pix) | 1.045 |
| Flux (e<sup>-</sup>/pix/sec) | 15.5 |
| Frames per exposure | 20 |
| Exposure (e<sup>-</sup>/Å<sup>2</sup>) | 42.58 |
| Defocus range (μm) | 1.09 - 2.5 |
| Micrographs collected | 6,606 |
| Particles extracted/final | 1016378/59972 |
| Symmetry imposed | none (C1) |
| Map sharpening B-factor | 122.3 |
| Unmasked resolution at 0.143 FSC (Å) | 4.4 |
| Masked resolution at 0.143 FSC (Å) | 3.35 |

**Table S2:** Statistical validation for integrative modeling of protein complexes.

| Protein complex | Cluster precision | Cluster size | % XLs satisfied | CCC |
| --- | --- | --- | --- | --- |
| Ank1-open | 41.739 Å | 6,672 | 87% | 0.9785 |
| Ank1-closed | 43.884 Å | 4,452 | 95% | 0.9718 |
| Ank1-open w/<br>enzymes | 52.747 Å | 4,361 | 90% | 0.9921 |
| Ank1-closed w/<br>enzymes | 52.849 Å | 2,506 | 98% | 0.9442 |
| band<br>4.1-spectrin | 25.937 Å | 4,237 | 95% | 0.9866 |

**Table S3:**
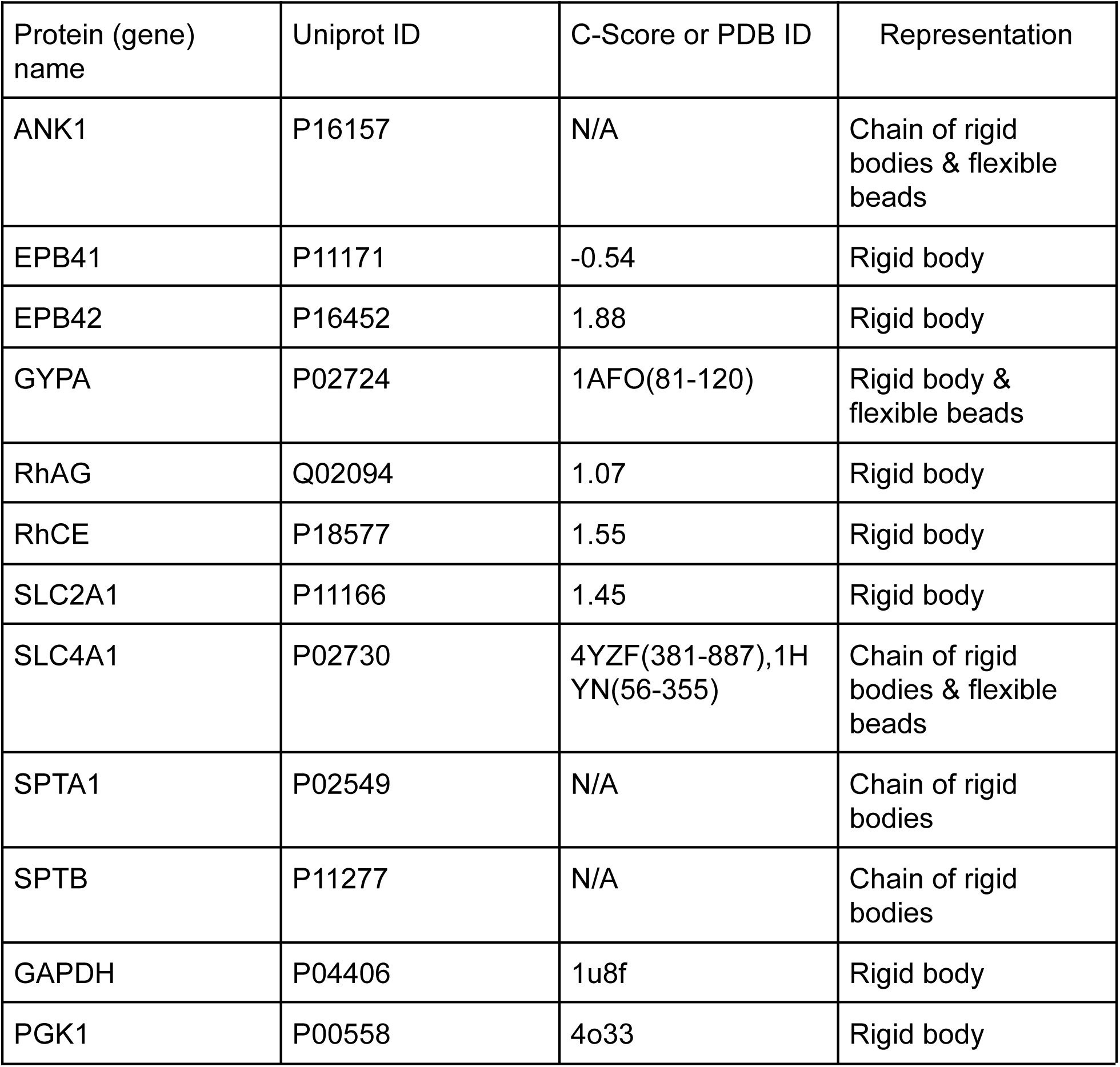
Individual protein modeling source and representation.

| Protein (gene) name | Uniprot ID | C-Score or PDB ID | Representation |
| --- | --- | --- | --- |
| ANK1 | P16157 | N/A | Chain of rigid bodies & flexible beads |
| EPB41 | P11171 | -0.54 | Rigid body |
| EPB42 | P16452 | 1.88 | Rigid body |
| GYPA | P02724 | 1AFO(81-120) | Rigid body & flexible beads |
| RhAG | Q02094 | 1.07 | Rigid body |
| RhCE | P18577 | 1.55 | Rigid body |
| SLC2A1 | P11166 | 1.45 | Rigid body |
| SLC4A1 | P02730 | 4YZF(381-887),1HYN(56-355) | Chain of rigid bodies & flexible beads |
| SPTA1 | P02549 | N/A | Chain of rigid bodies |
| SPTB | P11277 | N/A | Chain of rigid bodies |
| GAPDH | P04406 | 1u8f | Rigid body |
| PGK1 | P00558 | 4o33 | Rigid body |

